# Saruparib recapitulates PARP–BRCA synthetic lethality with PARP1-trapping properties limiting its cytotoxicity

**DOI:** 10.64898/2026.11.17.767429

**Authors:** Shiho Nakano, Hiroshi Onji, Ryo Kamata, Erina Kusakabe, Kaho Utsunomiya, Tomohiro Iguchi, Sho Isoyama, Shingo Dan, Kouji Hirota, Hiroyuki Sasanuma, Takashi Sugiyama, Yves Pommier, Kenjiro Sawada, Junko Murai

## Abstract

The original concept of PARP synthetic lethality proposes that loss of poly(ADP-ribose) polymerase (PARP) activity selectively kills homologous recombination (HR)-deficient cells. However, clinically approved PARP1/2 inhibitors (PARPis) are used at exposures exceeding those required for catalytic inhibition. At these doses, PARP trapping is cytotoxic to HR-proficient cells and limits the therapeutic window in HR-deficient cancers. Saruparib has been developed as a PARP1-selective inhibitor to reduce PARP2-associated toxicity; yet its PARP1-trapping properties remain incompletely characterized. Here, we show that saruparib exhibits trapping properties distinct from those of conventional PARPis. It causes minimal cytotoxicity in HR-proficient cells and remains minimally toxic in XRCC1-deficient cells, which are highly sensitive to trapping-associated cytotoxicity induced by conventional PARPis. Although methyl methanesulfonate, which induces extensive base damage, increases saruparib-induced PARP1 trapping and cytotoxicity, the overall trapping-associated cytotoxic response is weaker than that induced by olaparib and shows little further dose-dependent increase once catalytic PARP1 inhibition is achieved. Biochemical measurements demonstrated that PARP1–DNA complexes formed in the presence of saruparib are less stable than those in the presence of olaparib or talazoparib. Notably, saruparib potently kills BRCA1-, BRCA2-, and PALB2-deficient cells at concentrations sufficient for near-maximal catalytic inhibition, without a corresponding increase in cytotoxicity in HR-proficient cells. Thus, saruparib combines potent HR-deficient-selective cytotoxicity with limited PARP1 trapping and associated toxicity. Consequently, saruparib closely recapitulates the PARP–BRCA synthetic lethality while sparing HR-proficient cells, providing a mechanistic framework for its rational clinical use.

## Introduction

Poly(ADP-ribose) polymerase (PARP) inhibitors (PARPis) emerged as a new class of targeted anticancer agents following two landmark studies in 2005 demonstrating synthetic lethality between PARP inhibition and BRCA deficiency (1,2). These studies showed that genetic depletion of PARP1 or pharmacological inhibition of PARP activity was profoundly cytotoxic to BRCA-deficient cells, establishing the concept for the essentiality of PARP1 when homologous recombination-dependent DNA repair (HR) is compromised. At the time, this synthetic lethality was attributed primarily to the persistence of endogenous DNA lesions, which were converted during DNA replication into double-strand breaks requiring HR for repair. This discovery provided a compelling therapeutic concept: inhibition of a DNA repair enzyme could selectively kill BRCA-deficient tumor cells while largely sparing HR-proficient normal cells (3).

This concept led to the development of multiple clinical PARPis, including olaparib, rucaparib, niraparib, talazoparib, and veliparib. Although these inhibitors all potently suppress PARP catalytic activity in the low-nanomolar range, their cellular cytotoxicity differs by several orders of magnitude in both BRCA-deficient and HR-proficient cells (4). This discrepancy was subsequently explained by our earlier work identifying PARP trapping, whereby PARPis stabilize PARP1/2-DNA complexes that interfere with DNA replication and produce DNA damage (5,6). PARP-trapping potency varies markedly among clinical PARPis and parallels their cytotoxic potency and clinically tolerated doses, with the potent trapper talazoparib administered at substantially lower doses than relatively weaker trappers (7,8). However, this mechanism complicated the original concept of PARP–BRCA synthetic lethality. Unlike catalytic PARP inhibition, trapped PARP–DNA complexes themselves constitute highly cytotoxic replication-associated lesions that require dedicated processing mechanisms, including SPRTN-mediated proteolysis and p97-dependent extraction of PARP-DNA complexes from chromatin (5,9,10). Moreover, PARP trapping can also occur within DNA intermediates generated during Okazaki fragment processing behind replication forks, where persistent PARP–DNA complexes can promote ssDNA gap formation and ultimately cell death (11–13). Consequently, increasing PARP-trapping potency enhances cytotoxicity not only in HR-deficient cells but also in HR-proficient cells, including normal proliferating cells, thereby potentially limiting the therapeutic window of conventional PARPis.

The highly selective PARP1 inhibitor saruparib (AZD5305) was recently developed based on the hypothesis that selective inhibition and trapping of PARP1 while sparing PARP2 would retain antitumor efficacy while reducing the hematologic toxicity associated with conventional PARP1/2 inhibitors (14). This strategy was supported by evidence implicating PARP2 inhibition and/or trapping in PARPi-associated hematologic toxicity, particularly anemia (15,16). Accordingly, saruparib exhibits >500-fold selectivity for PARP1 over PARP2 and potent activity in BRCA1-, BRCA2-, PALB2-, and RAD51C-deficient models, with minimal effects on HR-proficient cells (14). Building on saruparib, AZD9574 was subsequently developed to retain potent and selective PARP1 inhibition while incorporating improved blood–brain barrier penetration (17).

Saruparib was initially characterized as a potent and selective PARP1 trapper using a cellular chromatin-retention assay in the presence of methyl methanesulfonate (MMS), a DNA-alkylating agent that induces extensive base damage (14). However, subsequent studies comparing saruparib with conventional PARPis in the absence of exogenous DNA-damaging agents reported substantially weaker PARP1 trapping by saruparib than that induced by olaparib or talazoparib (18,19). Genetic studies further revealed drug-specific allosteric regulation of PARP1–DNA interactions. The PARP1 L777P mutation, located in the regulatory helical domain and associated with increased conformational dynamics, shifts the response to saruparib toward PARP1 release from chromatin, whereas olaparib further enhances PARP1 trapping (20). Additionally, a recent study independently demonstrated saruparib-induced detrapping of several PARP1 catalytic mutants (21). Together, these apparently divergent observations suggest that saruparib modulates PARP1–DNA interactions differently from the currently clinically-approved PARPis and justifies further studies of its trapping properties.

In this study, we systematically investigated the pharmacological properties of saruparib across multiple genetic and DNA damage settings. We show that saruparib exhibits PARP1-trapping properties distinct from those of conventional PARPis, both with respect to the stability of trapped PARP1–DNA complexes and trapping-associated cytotoxicity. We conclude that saruparib’s pharmacological profile matches the original concept of PARP–BRCA synthetic lethality with potent cytotoxicity in HR-deficient cells, while exerting limited PARP trapping and cytotoxicity toward HR-proficient cells.

## Materials and Methods

### Cell lines and culture conditions

TOV112D (RRID: CVCL_3612) cells were obtained from the American Type Culture Collection (ATCC; Manassas, VA, USA). MCF-7 (RRID: CVCL_0031) and TK6 (IVGT) (RRID: CVCL_AR68) cells were obtained from the JCRB Cell Bank (https://cellbank.nibiohn.go.jp/english/). MCF-7 XRCC1-KO cells, derived from MCF-7 cells (RRID: CVCL_0031), were generated as previously described (22). TK6 PARP1-KO (22), XRCC1-KO (23) cells were generated as previously described. CCRF-CEM (RRID: CVCL_0207) cells were obtained from the Developmental Therapeutics Program (DTP), National Cancer Institute (NCI). Tree92 (RRID: CVCL_M659) cells were kindly provided by the Department of Hematology, Graduate School of Medicine, Kyoto University. SLFN11-KO CCRF-CEM and Tree92 cells were generated as previously described (24,25). Capan-1 (RRID: CVCL_0237) and PEO1 (RRID: CVCL_2686) parental cells and their BRCA2-revertant derivatives were kindly provided by Dr. Taniguchi (26,27). TK6 cells carrying auxin-inducible degron (AID)-tagged BRCA1, BRCA2, or PALB2 were generated as previously described (28,29).

TOV112D (RRID: CVCL_3612), MCF-7 (RRID: CVCL_0031), Capan-1 (RRID: CVCL_0237), and PEO1 (RRID: CVCL_2686) cells were cultured in Dulbecco’s modified Eagle medium (DMEM; FUJIFILM Wako Pure Chemical Industries, #044-29765) supplemented with 10% fetal bovine serum (FBS; Gibco, Thermo Fisher Scientific, #A5256701) and 1% antibiotic-antimycotic solution (100×; Nacalai Tesque, #09366-44). TK6 (IVGT; RRID: CVCL_AR68), CCRF-CEM (RRID: CVCL_0207), and Tree92 (RRID: CVCL_M659) cells were cultured in RPMI 1640 medium (FUJIFILM Wako Pure Chemical Industries, #189-02145) supplemented with 10% FBS (Gibco, Thermo Fisher Scientific, #A5256701) and 1% antibiotic-antimycotic solution (100×; Nacalai Tesque, #09366-44). Cells were cultured at 37°C in a humidified incubator with 5% CO₂. All cell lines were routinely tested for mycoplasma contamination and confirmed to be negative.

### Drug treatment

Saruparib (#S9875), niraparib (#S2741), veliparib (#S1004), and AZD9574 (#E2147) were obtained from Selleckchem. Talazoparib (#HY-16106) and the PARG inhibitor PDD00017273 (#HY-108360) were obtained from MedChemExpress. Olaparib was obtained from LC Laboratories (#O-9201). Methyl methanesulfonate (MMS) was obtained from Sigma-Aldrich (#129925-5G). 3-Indoleacetic acid (heteroauxin) was obtained from Nacalai Tesque (#19119-61).

### Auxin-inducible degron (AID)-mediated protein depletion

BRCA1, BRCA2, and PALB2 were conditionally depleted in TK6 cells using an auxin-inducible degron (AID) system (30). Cells were treated with 500 µM auxin (indole-3-acetic acid) to induce degradation of the AID-tagged proteins. In all experiments, auxin was added simultaneously with the indicated drugs and maintained throughout the treatment period. Efficient depletion of the AID-tagged proteins was confirmed within 6 h after auxin addition.

### Antibodies

Antibodies against BRCA1 (#9010, RRID: AB_2228244), BRCA2 (#10741, RRID: AB_2797730), PAR (#89190, RRID: AB_3716623), XRCC1 (#2735, RRID: AB_2218471), PARP1 (#9532, RRID: AB_659884), cleaved PARP (Asp214) (#5625, RRID: AB_10699459), RPA32/RPA2 (E8X5P) (total RPA2) (#35869, RRID: AB_2799086), cleaved Caspase3 (Asp175) (5A1E) (#9664, RRID: AB_2070042), histone H3 (#4499, RRID: AB_10544537), and GAPDH (#97166, RRID: AB_2756824) were obtained from Cell Signaling Technology; antibodies against MCM2 (#10513-1-AP, RRID: AB_2142131) and PARP1 (#13371-1-AP, RRID: AB_2160459; for PARP trapping assays) were obtained from Proteintech; antibody against α-tubulin (#017-25031, RRID: AB_2943605) was obtained from FUJIFILM Wako Pure Chemical Corporation; antibody against phospho-histone H2AX (Ser139) (#05-636, RRID: AB_309864) was obtained from Merck; antibody against PALB2 (#A301-246A, RRID: AB_890607) was obtained from Bethyl Laboratories.

### Immunoblotting

Whole-cell lysates were prepared by lysing cells in Laemmli SDS sample buffer followed by boiling. Protein concentrations were determined using an RC DC Protein Assay Kit (Bio-Rad Laboratories, #50000119). Immunoblotting was performed using standard procedures. The primary antibodies described in the Antibodies section above were used at a dilution of 1:3000 for PAR, XRCC1, PARP1, cleaved PARP, cleaved Caspase3, MCM2, BRCA1, BRCA2, and PALB2, GAPDH, α-tubulin; 1:5000 for histone H3; and 1:10000 for γH2AX. Membranes were then incubated with horseradish peroxidase–conjugated secondary antibodies, including anti-rabbit IgG (Promega, #W401B, RRID: AB_430833) and anti-mouse IgG (Promega, #W402B, RRID: AB_430834). Protein signals were detected using an iBright CL1500 Imaging System (Invitrogen).

Chromatin fractionation was performed using a commercial chromatin isolation kit, Subcellular Protein Fractionation Kit for Cultured Cells (Thermo Scientific, #78840), according to the manufacturer’s protocol, followed by immunoblotting as described above.

### Small interfering RNA (siRNA) transfection

TOV112D cells were transfected with SMARTpool ON-TARGETplus Human XRCC1 siRNA (Horizon Discovery, #L-009394-00-0005), Human BRCA2 siRNA (Horizon Discovery, #L-003462-00-0010), Human BRCA1 siRNA (Horizon Discovery, # L-003461-00-0005), Human PALB2 siRNA (Horizon Discovery, #L-012928-01-0005), and ON-TARGETplus Non-targeting Control Pool (Horizon Discovery, #D-001810-10-05) as a negative control. siRNAs (10 nM) were transfected using Lipofectamine RNAiMAX Reagent (Invitrogen, #13778-150) according to the manufacturer’s instructions. The culture medium was replaced 6 h after transfection, and subsequent experiments were performed 24 h after transfection. Knockdown efficiency was confirmed by immunoblotting.

### Cell viability assays

TOV112D and TK6 cells were seeded in 96-well white plates (SPL Life Sciences, #s30196) at a density of 3,000–10,000 cells per well in 100 μL of medium. Cells were continuously exposed to the indicated drug concentrations for the indicated time periods, with three technical replicate wells per condition. Experiments were independently repeated two or three times (biological replicates), as indicated in the corresponding figure legends. Cell viability was measured using an ATPlite 1step Luminescence ATP Detection Assay System (Revvity, #6016731) according to the manufacturer’s instructions. Luminescence was measured using a Synergy LX Multi-Mode Reader (BioTek). Cell viability (%) was normalized to untreated controls.

### Immunofluorescence analysis

TK6 and CCRF-CEM cells were cytocentrifuged onto SuperFrost Plus microscope slides (Thermo Fisher Scientific, #J1820AMNZ) using a Cytospin 4 Cytocentrifuge (Thermo Fisher Scientific). Cells were fixed with 4% paraformaldehyde in PBS for 15 min. Permeabilization was performed with 0.1% Triton X-100 in PBS for 15 min. Cells were blocked with 4% bovine serum albumin (BSA) in PBST for 15 min, followed by overnight incubation at 4 °C with anti-PAR antibody (#89190, RRID: AB_3716623; 1:300) and anti-phospho-histone H2AX (Ser139) antibody (#05-636, RRID: AB_309864; 1:300) diluted in blocking buffer. After washing, cells were incubated with appropriate secondary antibodies, including goat anti-rabbit IgG (H+L) cross-adsorbed Alexa Fluor 488 (Invitrogen, #A-11008, RRID: AB_143165) and goat anti-mouse IgG (H+L) cross-adsorbed Alexa Fluor 488 (Invitrogen, #A-11001, RRID: AB_2534069), at a dilution of 1:1000 in 3– 5% BSA/PBST for 1h at room temperature. After washing, cells were mounted with Fluoroshield Mounting Medium with DAPI (Abcam, #ab104139). Images were acquired using a BZ-X1000 fluorescence microscope (KEYENCE).

### PARP trapping assay by immunofluorescence

PARP1 trapping was assessed by immunofluorescence using a sequential extraction and fixation procedure adapted from Petropoulos et al. (19). TK6 cells were cytocentrifuged onto SuperFrost Plus microscope slides (Thermo Fisher Scientific, #J1820AMNZ) using a Cytospin 4 Cytocentrifuge (Thermo Fisher Scientific), whereas TOV112D and MCF7 cells were grown on chamber slides (Watson, #192-004). Following the indicated treatments, cells were first pre-extracted for 2 min with ice-cold 1× PBS containing 0.2% Triton X-100 and then fixed with 4% formaldehyde for 10 min at room temperature. After three washes with 1× PBS, cells were permeabilized with 1× PBS containing 0.2% Triton X-100 for 15 min at room temperature. The cells were then further extracted with ice-cold cytoskeleton buffer (0.5% Triton X-100, 10 mM PIPES, pH 6.8, 3 mM MgCl2, 200 mM NaCl, and 300 mM sucrose) for 10 min at 4 °C, followed by fixation with ice-cold methanol for 15 min at −20 °C. Following three washes with PBS, cells were blocked with 5% BSA/PBS for 1 h at room temperature and incubated overnight at 4 °C with a PARP1 rabbit polyclonal antibody (1:1,000; Proteintech, #13371-1-AP, RRID: AB_2160459) diluted in 5% BSA/PBS. After three washes with PBS, cells were incubated with the appropriate secondary antibody diluted in 5% BSA/PBS for 1 h at room temperature in the dark. For TOV112D and MCF7 cells, Hoechst 33342 (Invitrogen, #H3570) was added at a dilution of 1:2,000 during secondary antibody incubation for nuclear staining. For TK6 cells, slides were mounted with Fluoroshield Mounting Medium with DAPI (Abcam, #ab104139). Images were acquired using a BZ-X1000 fluorescence microscope (KEYENCE).

### Immunofluorescence analysis with or without EdU labeling

MCF-7 cells were grown on chamber slides (Watson, #192-004) and treated with the indicated drugs. To label cells undergoing DNA synthesis, cells were incubated with 10 μM 5-ethynyl-2′-deoxyuridine (EdU) for 1 h, pre-extracted with ice-cold PBS containing 0.2% Triton X-100 for 1 min, fixed with 4% paraformaldehyde in PBS for 15 min, and blocked with 5% BSA/PBS for 10 min. Next, for EdU detection, cells were incubated with Click-iT Plus reaction cocktail for 30 min (Invitrogen, #C10640). Cells were incubated overnight with primary antibodies in 3% BSA/PBS at 4 °C. After washing with TBST, the cells were incubated with proper secondary antibodies and Hoechst33342 in 3% BSA/TBST for 1 h. After washing with TBST, cells were mounted with ProLong Gold Antifade Mountant (Invitrogen, # P36930). Images were acquired using a BZ-X1000 fluorescence microscope (KEYENCE).

### Cell cycle analysis by flow cytometry

Cells were incubated with 10 μM EdU for 1 h using a Click-iT Plus EdU Alexa Fluor 488 Flow Cytometry Assay Kit (Invitrogen, #C10633) according to the manufacturer’s instructions. DNA content was assessed by propidium iodide (PI) staining (Nacalai Tesque, #29037-34). Data were acquired using a CytoFLEX flow cytometer (Beckman Coulter) and analyzed with FlowJo software (BD Biosciences, version 10.10.0, RRID:SCR_008520).

### Fractionation of PARP-DNA complexes

TK6 (IVGT) (RRID: CVCL_AR68) cells (3 x 10^6^) were treated with or without the indicated drugs for 4 h. Fractionation of PARP–DNA complexes using buffers of increasing stringency was performed as previously described (5).

### Preparation of nicked DNA substrates

Oligonucleotides were synthesized by Eurofins Genomics (Tokyo, Japan). A 30-nucleotide 5′-biotinylated oligonucleotide (5′-BioON-AGGATGACCTGATCGTACGATGCTAGTCGA-3′) was used as the common top strand. To prepare a DNA duplex containing a single-strand break with 5′-deoxyribose phosphate (5′-dRP) and 3′-OH termini, the biotinylated oligonucleotide was annealed with a complementary 30-mer containing a centrally positioned deoxyuridine (5′-TCGACTAGCATCGUACGATCAGGTCATCCT-3′). The two strands were mixed at equimolar concentrations in annealing buffer containing 10 mM Tris-HCl and 50 mM NaCl, heated at 95°C for 2 min, and gradually cooled to room temperature. The uracil-containing duplex was incubated with 5 U uracil-DNA glycosylase (UDG; New England Biolabs, #M0280S) for 10 min at 37°C, followed by incubation with 10 U APE1 (New England Biolabs, #M0282S) for 30 min at 37°C in 1× NEBuffer 4. This sequential treatment generated a centrally located nick containing 5′-dRP and 3′-OH termini. The DNA substrates were purified using an Oligo Clean C Concentrator kit (Zymo Research, D4060) and eluted in nuclease-free water. The enzyme concentrations are consistent with the NEB product specifications: M0280S is 5 U/μL and M0282S is 10 U/μL. For preparation of a duplex containing a 5′-phosphate/3′-OH nick, the 5′-biotinylated 30-mer was annealed with a 5′-phosphorylated 15-mer (5′-P-CGATCAGGTCATCCT-3′) and an adjacent 15-mer (5′-TCGACTAGCATCGTA-3′), both complementary to the biotinylated strand.

### Biolayer interferometry analysis of PARP1–DNA complex stability

PARP1–DNA binding and dissociation were analyzed by biolayer interferometry (BLI) using an Octet R4 instrument (Sartorius) equipped with Streptavidin (SA) Biosensors (Sartorius, 18-5019). Unless otherwise indicated, measurements were performed in Buffer A consisting of 50 mM Tris-HCl (pH 8.0), 4 mM MgCl₂, 0.1 mg/mL bovine serum albumin, and 0.05% Tween-20, supplemented with KCl or NaCl as specified. SA biosensors were equilibrated in Buffer A for 60 s and loaded with the 5′-biotinylated nicked DNA substrate (0.5 ng/μL) for 120 s. After a second 60-s equilibration in Buffer A, DNA-loaded biosensors were transferred to Buffer A containing 25 nM recombinant human PARP1 (Invitrogen, RP-88140) for a 300-s association step. During baseline, loading, and association, Buffer A contained 100 mM KCl or 100 mM NaCl, depending on the experimental condition. PARP1–DNA complexes were then transferred to protein-free Buffer A containing DMSO, 10 μM olaparib, 10 μM saruparib, or 1 μM talazoparib, in the absence or presence of 1 mM NAD⁺ (Selleck Chemicals, S2518), and dissociation was monitored for 300 s. During the dissociation step, the concentrations of KCl or NaCl were varied as indicated in the figures. For experiments using the 5′-dRP/3′-OH nicked DNA substrate, dissociation was examined in KCl- or NaCl-containing buffers. Additional experiments were performed using the 5′-phosphate/3′-OH nicked DNA substrate. In Fig. 4 and Fig. S5, dissociation was compared in buffers containing 100 or 150 mM KCl or NaCl, as indicated. Buffer-only sensors were included as references, and the mean signal from paired reference sensors was subtracted from the corresponding PARP1-containing sensor signals. No inter-step correction or Savitzky–Golay filtering was applied. Dissociation rate constants (k_dis) were determined using Octet Analysis Studio software (version 12.2.2.26) by local fitting of the 300-s dissociation phase to a 1:1 binding model without constraining the dissociation endpoint to return to the pre-association baseline.

### High-content imaging and single-cell analysis

TOV112D cells were seeded at 4,000 cells per well in 100 μL of culture medium in PhenoPlate 96-well microplates (Revvity, #6055302) and incubated overnight. Following siRNA transfection, cells were treated with the indicated drugs for 4 h and subsequently subjected to immunofluorescence staining as described above using the anti-phospho-histone H2AX (Ser139) antibody (#05-636, RRID: AB_309864) and an Alexa Fluor 488-conjugated secondary antibody. Hoechst was used for nuclear staining. After the final washing step, 100 μL PBS was added to each well, and the plates were subjected to imaging and single-cell analysis using an Operetta CLS high-content analysis system (Revvity, #HH16000020). Nuclei were identified based on Hoechst staining, and Hoechst sum intensity and Alexa 488 sum intensity were quantified for individual cells. Hoechst sum intensity was used as a measure of cellular DNA content, whereas Alexa 488 sum intensity was defined as the integrated fluorescence intensity of all γH2AX foci detected within each cell. Single-cell γH2AX intensity was then analyzed in relation to DNA content.

### Statistical analysis

Data are presented as mean ± standard deviation (SD). Details of technical and biological replicates are described in each figure legend. No formal power analysis was performed to determine sample size. No formal randomization was performed for the in vitro experiments. Investigators were not blinded to treatment conditions during data acquisition or analysis. Statistical analyses were performed using GraphPad Prism (GraphPad Software, version 11.1.0, RRID: SCR_002798). For comparisons among more than two groups, one-way ANOVA followed by Tukey’s multiple-comparison test was used. *P* values < 0.05 were considered statistically significant. Significance is indicated as follows: NS, not significant; \**P* < 0.05; \*\**P* < 0.01; \*\*\**P* < 0.001; \*\*\*\**P* < 0.0001.

### Data and Code availability

The source data underlying the findings of this study have been deposited in Mendeley Data (DOI: 10.17632/rxmxxhr49v.1). The dataset is currently available to reviewers through the following private access link: https://data.mendeley.com/preview/rxmxxhr49v?a=34ae9a58-e678-41ea-82d0-4bc0afe1c3f3

The dataset will be made publicly available upon acceptance of the manuscript.

## Results

### Saruparib potently inhibits PARP1 catalytic activity with limited PARP1-dependent cytotoxicity

PARP1 trapping contributes substantially to the cytotoxicity of conventional PARP inhibitors (PARPis), as evidenced by the reduced sensitivity of PARP1-knockout (PARP1-KO) cells compared with their wild-type (WT) counterparts (5,31). To compare PARP1-dependent cytotoxicity among PARPis, we examined human B-lymphoblastoid TK6 WT and PARP1-KO cells (Fig. 1A and Fig. S1A). WT cells were sensitive to talazoparib, niraparib and olaparib whereas sensitivity was markedly attenuated in PARP1-KO cells. In contrast, saruparib and AZD9574 showed minimal cytotoxicity in either WT or PARP1-KO cells, even at high concentrations (25–50 μM). Thus, unlike conventional PARPis, saruparib and AZD9574 induce minimal PARP1-dependent cytotoxicity under unperturbed conditions.

**Figure 1.**
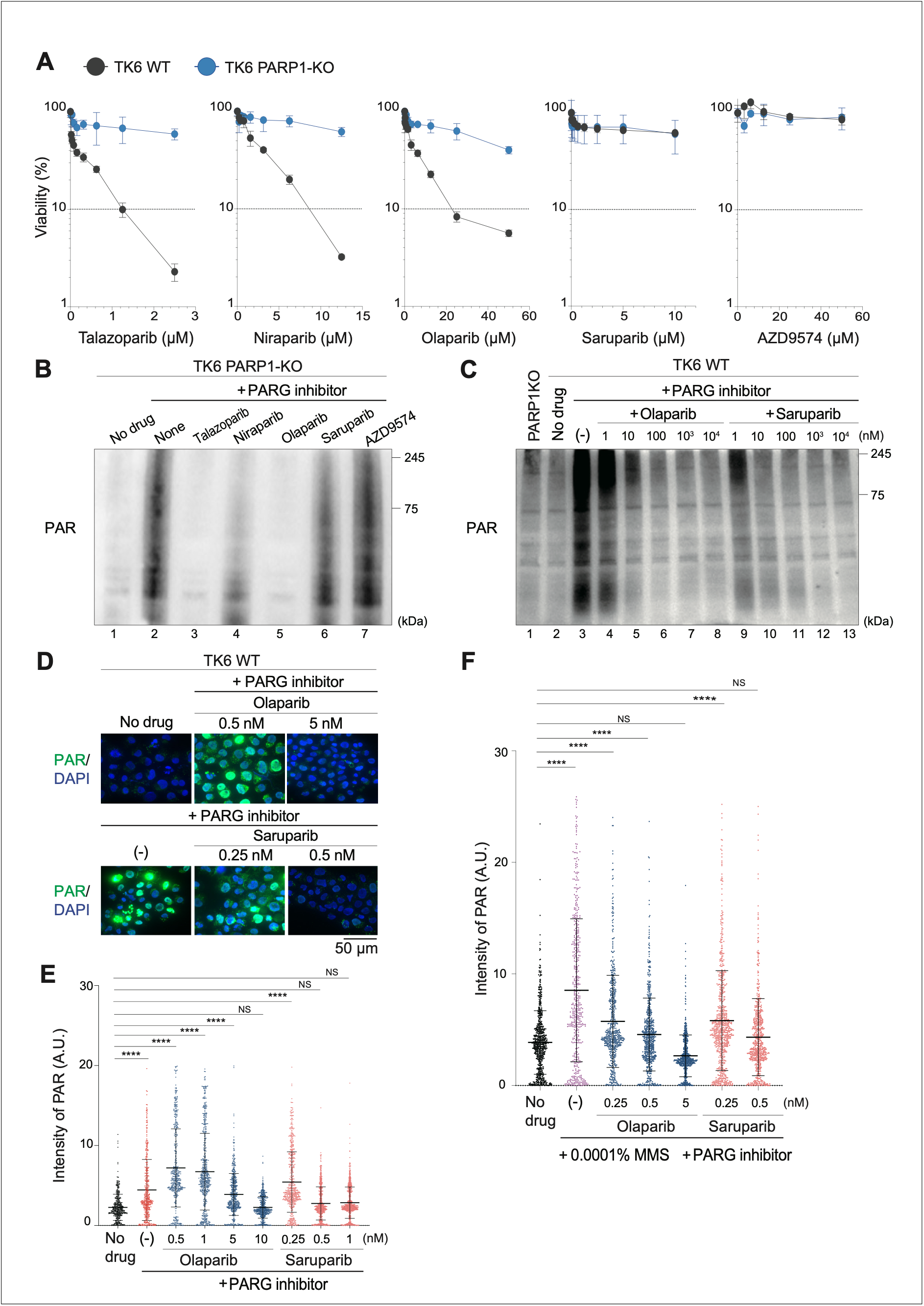
Saruparib potently inhibits PARP1 catalytic activity with limited PARP1-dependent cytotoxicity. A, Cell viability of TK6 WT and PARP1-KO cells treated with the indicated PARP inhibitors for 48 h. Viability was normalized to untreated controls. Data are mean ± SD of triplicate wells and are representative of two independent experiments. **B,** Immunoblot analysis of PAR in TK6 PARP1-KO cells treated with the indicated PARP inhibitors (100 nM) in the presence of PARG inhibitor (10 μM) for 12 h. Cells treated with PARG inhibitor alone and untreated cells without PARG inhibitor were included as controls. **C,** Dose-dependent inhibition of PAR formation by olaparib or saruparib in TK6 WT cells. Cells were treated with the indicated concentrations of olaparib or saruparib (1–10,000 nM) in the presence of PARG inhibitor (10 μM) for 12 h. PARP1-KO cells, untreated TK6 WT cells, and TK6 WT cells treated with PARG inhibitor alone were included as controls. **D,** Representative immunofluorescence images of PAR (green) and Hoechst-stained nuclei (blue) in TK6 WT cells treated with olaparib or saruparib at the indicated concentrations in the presence of PARG inhibitor (10 μM) for 4 h. Scale bar, 50 μm. **E,** Ǫuantification of PAR fluorescence intensity in **D**. Each dot represents an individual cell; bars indicate mean ± SD (n = 440–824 cells per condition). Data are representative of two independent experiments. **F,** PAR fluorescence intensity in TK6 WT cells treated with olaparib or saruparib at the indicated concentrations in the presence of MMS (0.0001%) and PARG inhibitor (10 μM) for 4 h. Data are presented as in **E** (n = 556–724 cells per condition). For **E** and **F**, NS, not significant; ****P < 0.0001 by one-way ANOVA followed by Tukey’s multiple-comparisons test.

We next confirmed the PARP1 selectivity of saruparib and AZD9574 by specifically assessing PARP1-independent PARylation in PARP1-KO cells treated with a poly(ADP-ribose) glycohydrolase inhibitor (PARGi) (Fig. 1B). In the absence of PARP1, PARG inhibition revealed a detectable residual PAR signal, reflecting predominantly PARP2-mediated PAR synthesis (Fig. 1B, lane 2). This residual PAR accumulation was strongly suppressed by talazoparib and olaparib and partially suppressed by niraparib, consistent with their inhibition of PARP2 catalytic activity. In contrast, saruparib and AZD9574 did not appreciably reduce PARylation under these conditions, supporting their selectivity for PARP1 over PARP2.

To examine whether the lack of PARP2 inhibition by saruparib affects its ability to suppress overall cellular PARylation, we next compared olaparib and saruparib in TK6 WT (PARP1/2-proficient) cells. Olaparib at 10 nM and saruparib at 1 nM produced comparable PAR suppression in PARGi-treated WT cells (Fig. 1C, lanes 5 and 9), whereas 100 nM olaparib and 10 nM saruparib almost completely suppressed PARylation (Fig. 1C, lanes 6 and 10). The absence of a substantial residual PAR signal with saruparib, despite its minimal inhibition of PARP2, further indicates that PARP1 is the predominant source of cellular PARylation under these conditions. Ǫuantitative immunofluorescence confirmed this approximately tenfold difference in potency, with 5 nM olaparib and 0.5 nM saruparib producing similar PAR suppression (Fig. 1D, E). Comparable results were obtained in human T-lymphoblast CCRF-CEM cells (Fig. S1B). Under conditions of robust PARP1 activation induced by MMS in the presence of PARGi, a similar difference was observed (Fig. 1F). Thus, saruparib inhibits cellular PARP1 catalytic activity with approximately tenfold greater potency than olaparib, even under conditions of enhanced PARP1 activation.

### Saruparib exhibits minimal PARP1 trapping even in XRCC1-deficient cells

XRCC1 facilitates single-strand break repair and prevents excessive PARP1 engagement at DNA breaks; consequently, XRCC1-deficient cells accumulate trapped PARP1–DNA complexes and are highly sensitive to PARP-trapping drugs (22,32,33). In TK6 cells, XRCC1 loss, confirmed by immunoblotting (Fig. S2A), markedly increased sensitivity to talazoparib, niraparib and olaparib, but not to saruparib or AZD9574 (Fig. 2A). We extended this analysis to MCF7 breast cancer and TOV112D ovarian cancer cells with XRCC1 knockout (XRCC1-KO) and knockdown (siXRCC1), respectively (Fig. S2A). XRCC1 loss markedly enhanced sensitivity to olaparib, whereas saruparib sensitivity was modestly increased in MCF7 cells and minimal in TOV112D cells (Fig. 2B, S2B). Notably, saruparib cytotoxicity reached a plateau in the low-nanomolar range, whereas olaparib cytotoxicity continued to increase across the micromolar range in both cell lines.

**Figure 2.**
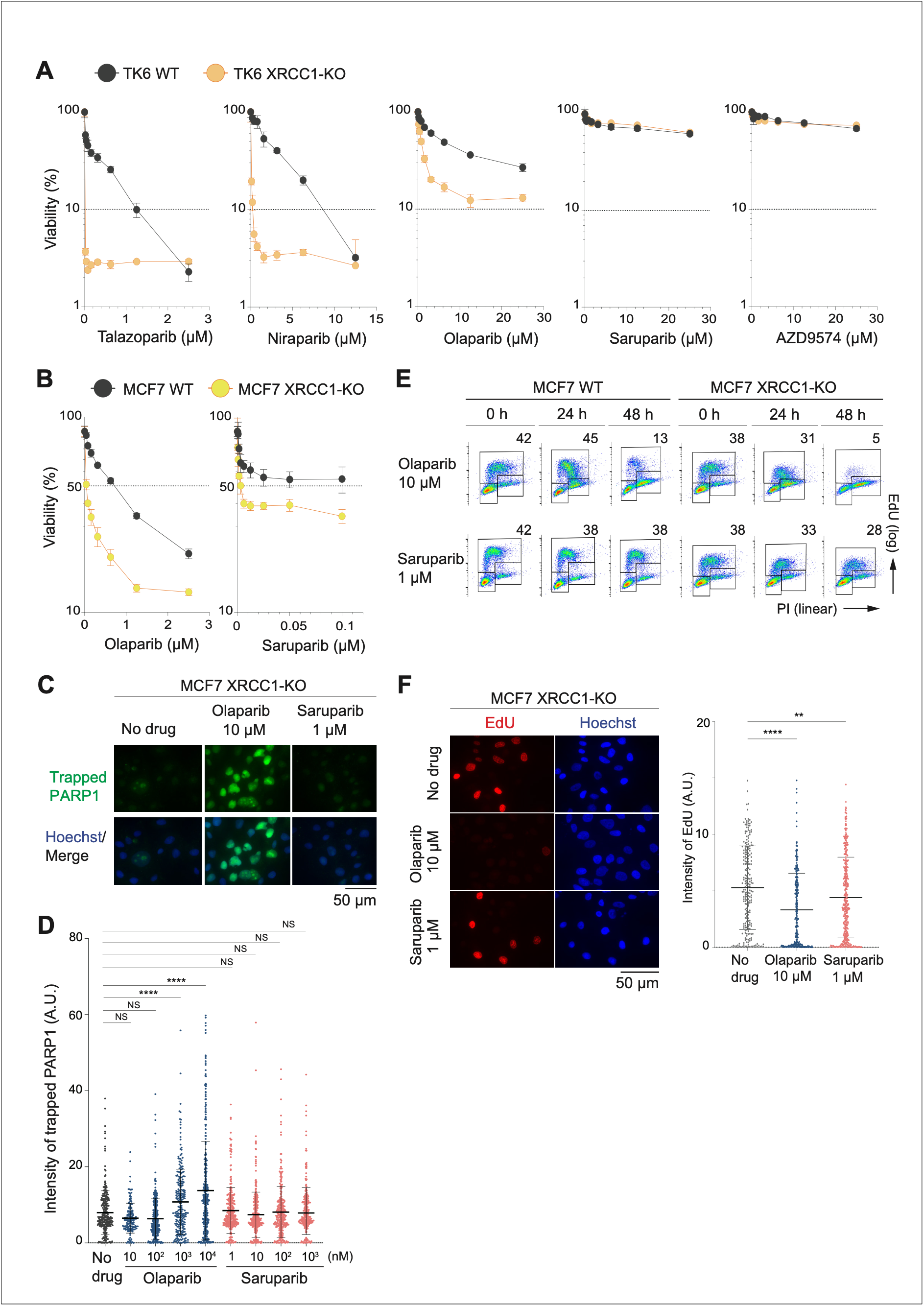
Saruparib exhibits minimal PARP1 trapping even in XRCC1-deficient cells. **A,** Cell viability of TK6 WT and XRCC1-KO cells treated with the indicated PARP inhibitors for 48 h. Viability was normalized to the corresponding untreated controls. Data are mean ± SD of triplicate wells and are representative of three independent experiments. **B,** Cell viability of MCF7 WT and XRCC1-KO cells treated with olaparib or saruparib for 6 days. Viability was normalized to the corresponding untreated controls. Data are mean ± SD of triplicate wells and are representative of three independent experiments. **C,** Representative immunofluorescence images of chromatin-bound PARP1 in MCF7 XRCC1-KO cells treated with olaparib (10 μM) or saruparib (1 μM) for 24h. Soluble proteins were removed by pre-extraction before fixation. Chromatin-bound PARP1 is shown in green and Hoechst-stained nuclei in blue. Scale bar, 50 μm. **D,** Ǫuantification of chromatin-bound PARP1 fluorescence intensity in MCF7 XRCC1-KO cells treated with the indicated concentrations of olaparib or saruparib. Each dot represents an individual cell; bars indicate mean ± SD (n = 241–361 cells per condition). Data are representative of two independent experiments. **E,** Cell-cycle distribution of MCF7 WT and XRCC1-KO cells treated with olaparib (10 μM) or saruparib (1 μM) for 24 or 48 h. Cells were analyzed by EdU incorporation and propidium iodide (PI) staining followed by flow cytometry. Numbers indicate the percentage of cells in the replicating S-phase population. Data are representative of two independent experiments. **F,** Representative immunofluorescence images and quantification of EdU incorporation in MCF7 XRCC1-KO cells treated with olaparib (10 μM) or saruparib (1 μM) for 24 h. Cells were pulse-labeled with EdU for 2 min before fixation. EdU is shown in red and Hoechst-stained nuclei in blue. Scale bar, 50 μm. Each dot represents an individual cell; bars indicate mean ± SD (n = 278–407 cells per condition). Data are representative of two independent experiments. For **D** and **F**, NS, not significant; **P < 0.01; ****P < 0.0001 by one-way ANOVA followed by Tukey’s multiple-comparisons test.

We next directly assessed chromatin-associated PARP1 using an immunofluorescence-based trapping assay with pre-extraction of soluble proteins in MCF7 XRCC1-KO and TOV112D siXRCC1 cells. Olaparib induced a concentration-dependent increase in chromatin-associated PARP1 at concentrations up to 10 μM, whereas saruparib showed no detectable PARP1 trapping at concentrations up to 1 μM (Fig. 2C, D, S2C, D).

Because trapped PARP1 can interfere with DNA replication, we next examined cell-cycle progression by EdU incorporation. Olaparib reduced the S-phase population, particularly in XRCC1-KO cells (Fig. 2E upper panels), whereas saruparib caused no apparent alteration in cell-cycle distribution in either WT or XRCC1-KO cells (Fig. 2E lower panels). Immunofluorescence analysis after pre-extraction combined with EdU labelling further confirmed replication suppression by olaparib but minimal replication suppression by saruparib (Fig. 2F). Together, these results demonstrate that saruparib has minimal PARP1-trapping activity even under conditions that strongly favor PARP1 trapping.

### Saruparib induces saturable PARP1 trapping in the presence of MMS

Because previous studies have reported PARP1 trapping by saruparib under conditions of exogenous DNA damage (14), we examined PARPi cytotoxicity in the presence of MMS. Olaparib-induced cytotoxicity progressively increased with both olaparib and MMS concentrations, extending into the micromolar range (Fig. 3A). In contrast, although saruparib markedly enhanced MMS-induced cytotoxicity, its effect reached a plateau at approximately 10 nM, with little further increase at higher concentrations (Fig. 3A). The maximal cytotoxicity of saruparib was generally lower than that of olaparib. Talazoparib and veliparib showed concentration-response profiles similar to that of olaparib, although over different concentration ranges (Fig. 3A). Thus, unlike conventional PARPis, saruparib-induced cytotoxicity rapidly saturates at low-nanomolar concentrations in the presence of MMS.

**Figure 3.**
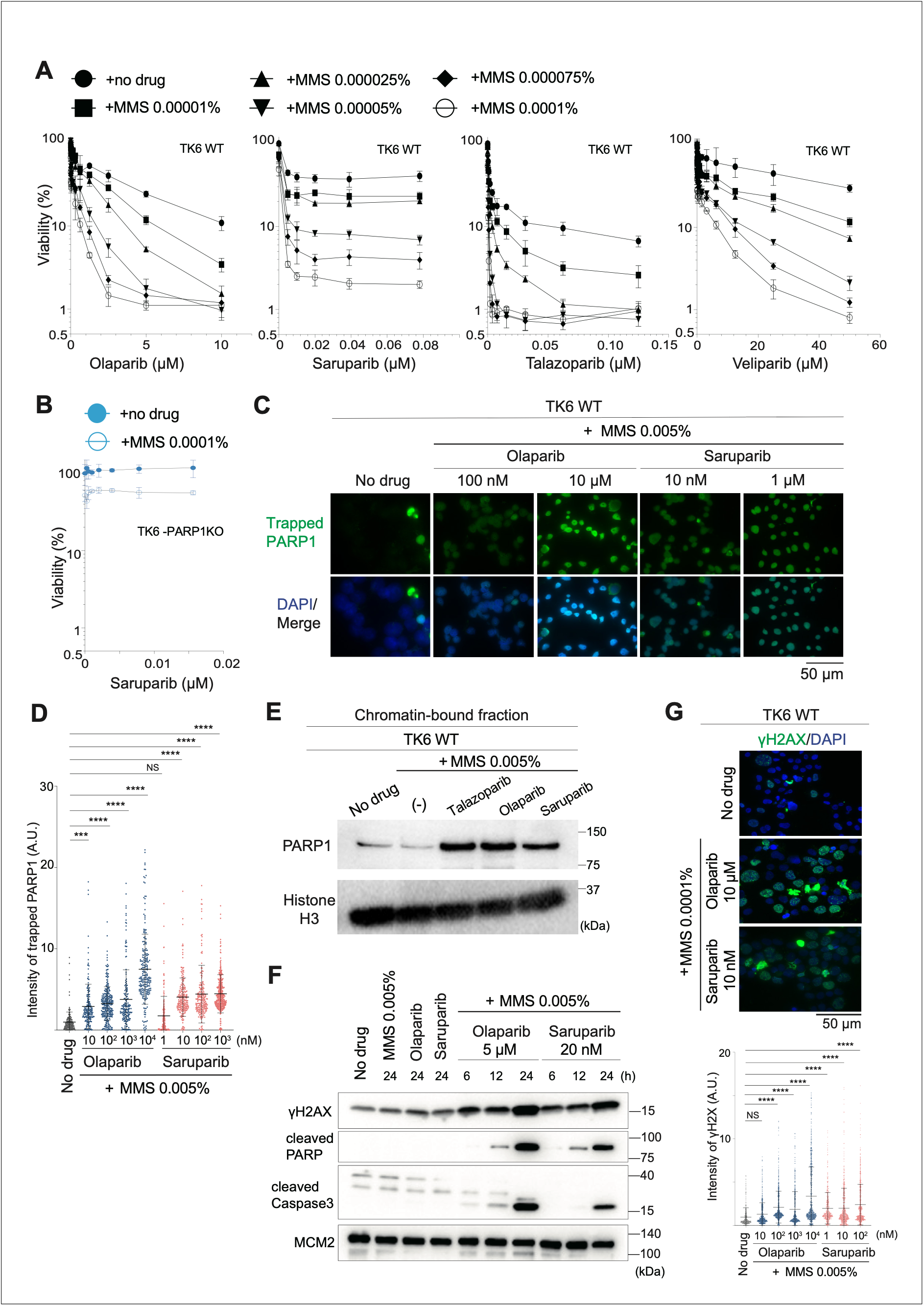
Saruparib induces saturable PARP1 trapping in the presence of MMS. **A,** Cell viability of TK6 WT cells treated with the indicated concentrations of PARP inhibitors alone or in combination with MMS for 48 h. Viability was normalized to untreated controls. Data are mean ± SD of triplicate wells and are representative of two or three independent experiments, depending on the inhibitor. **B,** Cell viability of TK6 PARP1-KO cells treated with saruparib alone or in combination with MMS (0.0001%) for 48 h. Data are presented as in **A**. **C,** Representative images of chromatin-bound PARP1 in TK6 WT cells treated with olaparib or saruparib in combination with MMS (0.005%), assessed as in Fig. 2C. Scale bar, 50 μm. **D,** Ǫuantification of chromatin-bound PARP1 fluorescence intensity in TK6 WT cells treated with the indicated concentrations of olaparib or saruparib in combination with MMS (0.005%). Each dot represents an individual cell; bars indicate mean ± SD (n = 184–389 cells per condition). Data are representative of three independent experiments. **E,** Immunoblot analysis of PARP1 in chromatin-bound fractions from TK6 WT cells treated with the indicated PARP inhibitors (10 μM) in combination with MMS (0.005%) for 4 hours. Histone H3 served as a loading control. **F,** Time-course immunoblot analysis of γH2AX, cleaved PARP, and cleaved caspase-3 in TK6 WT cells treated with olaparib or saruparib in combination with MMS (0.005%). MCM2 served as a loading control. Data are representative of three independent experiments. **G,** Representative images and quantification of γH2AX in TK6 WT cells treated with the indicated concentrations of olaparib or saruparib in combination with MMS (0.0001%). Each dot represents an individual cell; bars indicate mean ± SD (n = 373–795 cells per condition). Scale bar, 50 μm. For **D** and **G**, NS, not significant; ****P < 0.0001 by one-way ANOVA followed by Tukey’s multiple-comparisons test.

To determine whether catalytic PARP1 inhibition alone accounts for the MMS-dependent cytotoxicity of saruparib, we compared the effects of saruparib in WT cells with the intrinsic MMS sensitivity of PARP1-KO cells. Treatment with low dose MMS (0.0001%) reduced viability to approximately 70% in PARP1-KO cells (Fig. 3B), whereas saruparib plus MMS reduced WT cell viability to approximately 2% (Fig. 3A). These results indicate that pharmacological inhibition of PARP1 by saruparib in the presence of MMS is substantially more cytotoxic than genetic loss of PARP1 under the same MMS concentration, suggesting that catalytic PARP1 inhibition alone cannot account for saruparib-induced cytotoxicity.

We therefore directly assessed PARP1 trapping using a pre-extraction immunofluorescence assay in TK6 WT and TOV112D cells. As expected, olaparib increased chromatin-associated PARP1 in a concentration-dependent manner (Fig. 3C-D, S3A). Saruparib also induced PARP1 trapping in the presence of MMS but trapping rapidly plateaued at approximately 10 nM (Fig. 3C-D, S3A). Immunoblotting of chromatin-bound proteins independently confirmed PARP1 trapping by talazoparib, olaparib and saruparib in the presence of MMS (Fig. 3E). Treatment with 20 nM saruparib plus MMS induced PARP1 and caspase-3 cleavage and markedly increased γH2AX to levels comparable to those induced by 5 μM olaparib plus MMS (Fig. 3F), demonstrating robust apoptosis and DNA damage. Consistently, saruparib at 1 nM in the presence of MMS significantly increased γH2AX, and this increase remained largely constant at concentrations up to 100 nM (Fig. 3G). In contrast, olaparib at 10 nM did not significantly increase γH2AX, whereas at 100 nM and above, it induced γH2AX roughly in a dose-dependent manner (Fig. 3G). Thus, saruparib induces PARP1 trapping under MMS-treated conditions, but this trapping is highly saturable and distinct from the progressively increasing trapping observed with conventional PARPis.

### Saruparib induces replication stress in the presence of MMS

As an additional approach to determine whether saruparib induces replication stress in the presence of MMS, we employed SLFN11-dependent assays (34–36). SLFN11 is a replication stress sensitizer that promotes cell death by blocking DNA replication through binding to single-stranded DNA (ssDNA) (24,37,38), together with multiple downstream pathways (39–41). Furthermore, SLFN11 sensitizes cancer cells to PARP inhibitors by promoting ssDNA gap formation behind replication forks (8). Consistent with our previous findings (42), olaparib showed selective cytotoxicity in SLFN11-positive parental CCRF-CEM and Tree92 cells compared with their SLFN11-KO counterparts (Fig. S4A, B). In contrast, saruparib showed minimal SLFN11-dependent cytotoxicity as a single agent (Fig. S4A, B).

We next examined the effects of PARP inhibition in the presence of increasing concentrations of MMS. In SLFN11-positive parental cells, MMS markedly potentiated the cytotoxicity of both olaparib and saruparib (Fig. S4C, D). Notably, saruparib-induced cytotoxicity reached a plateau at low-nanomolar concentrations, whereas olaparib cytotoxicity continued to increase with increasing drug concentrations. In contrast, MMS-dependent sensitization to either PARPi was markedly attenuated in SLFN11-KO cells (Fig. S4E, F). These results indicate that the cytotoxicity of saruparib plus MMS is strongly dependent on SLFN11, consistent with the induction of replication stress under these conditions.

To directly examine DNA replication, we analyzed EdU incorporation in Tree92 parental cells overexpressing the anti-apoptotic protein MCL1 to minimize apoptosis (25). MMS or PARPis alone had only a modest effect on DNA replication, whereas the combination of MMS with either olaparib or saruparib markedly reduced EdU incorporation (Fig. S4G). Saruparib produced pronounced replication perturbation at concentrations as low as 10 nM, with little further effect at 1 μM, whereas the effect of olaparib was substantially greater at 1 μM than at 10 nM. Thus, consistent with its saturable PARP1-trapping activity, saruparib induces robust replication stress in the presence of MMS that saturates at low-nanomolar concentrations.

### PARP1–DNA complexes formed in the presence of saruparib are less stable than those formed with the currently clinically-approved PARP inhibitors

To clarify the differential trapping modes of conventional PARPis and saruparib, we first assessed the stability of chromatin-bound PARP1 by sequential extraction with increasing concentrations of KCl (Fig. 4A, B). PARP1 trapped by olaparib remained associated with chromatin even under the highest-stringency extraction condition (fraction E). In contrast, PARP1 trapped in the presence of saruparib appeared in earlier, lower-stringency fractions (fractions C and D), suggesting less stable chromatin association.

**Figure 4.**
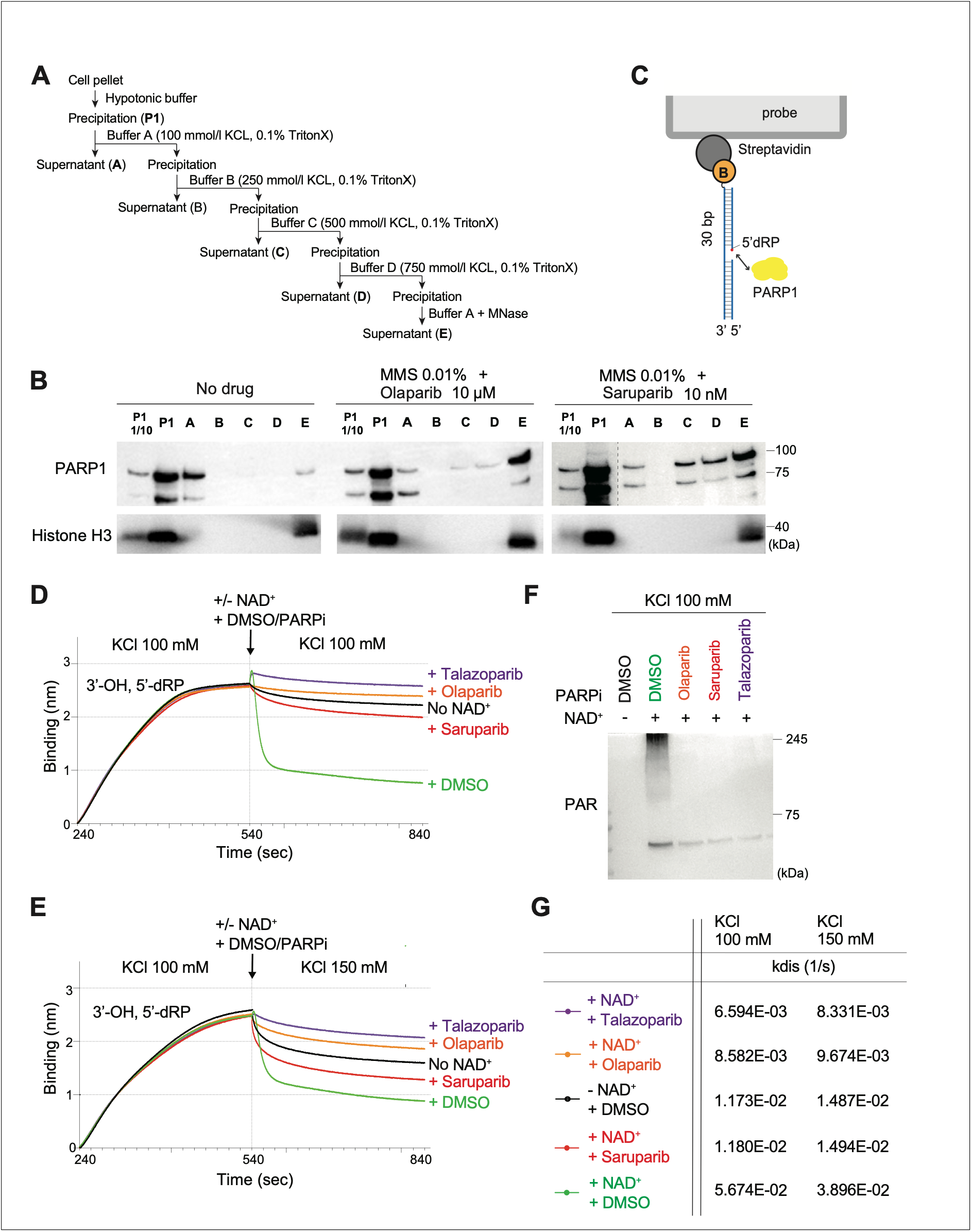
Saruparib forms less stable PARP1–DNA complexes than conventional PARP inhibitors. **A, B,** Sequential fractionation analysis of PARP1–DNA complexes in TK6 cells treated with MMS (0.01%) together with olaparib (10 μM) or saruparib (10 nM) for 4 h. **A,** Scheme for preparation of the indicated fractions (P1 and A–E) using buffers of increasing stringency. **B,** PARP1 and histone H3 in each fraction were analyzed by immunoblotting. One-tenth of the P1 fraction was loaded as indicated. **C,** Schematic of the biolayer interferometry (BLI) assay. A 30-bp biotinylated DNA probe containing a 3′-OH/5′-deoxyribose phosphate (5′-dRP) nick was immobilized on a streptavidin-coated biosensor for PARP1 binding. **D,** Representative BLI sensorgrams showing PARP1 dissociation from the 3′-OH/5′-dRP DNA substrate. Recombinant PARP1 (25 nM) was allowed to bind DNA-loaded biosensors for 300 s in Buffer A containing 100 mM KCl. Sensors were then transferred to protein-free buffer containing DMSO, olaparib (10 μM), saruparib (10 μM), or talazoparib (1 μM), with or without NAD⁺ (1 mM), and dissociation was monitored for 300 s. **E,** BLI analysis performed as in **D**, except that dissociation was monitored in 150 mM KCl. **F,** Dissociation solutions from the BLI assay in **D** were analyzed by immunoblotting for PAR to confirm PARP1 catalytic activity and its inhibition by the indicated PARP inhibitors. **G,** Dissociation rate constants (k_dis) calculated from the BLI curves in **D** and **E**.

To directly characterize the stability and dissociation kinetics of PARP1–DNA complexes, we established a cell-free biolayer interferometry assay using the Octet platform (Fig. 4C). Recombinant PARP1 was bound to a 30-bp double-stranded DNA substrate containing a centrally located single-strand break with 5′-deoxyribose phosphate (5′-dRP) and 3′-OH termini, and PARP1 dissociation was monitored in real time (Fig. 4D-G). In the absence of NAD⁺, PARP1 remained stably associated with DNA, providing a baseline for PARP1–DNA binding in the absence of catalytic activity, whereas NAD⁺ induced rapid dissociation, consistent with auto-PARylation-dependent release. In the presence of NAD⁺, olaparib and talazoparib slowed PARP1 dissociation even beyond the NAD⁺-free baseline, indicating stabilization of the PARP1–DNA complex beyond that resulting from loss of catalytic activity alone. In contrast, saruparib showed dissociation kinetics similar to the control without NAD⁺.

Under alternative experimental conditions using NaCl-containing buffers and a DNA substrate carrying a 5′-phosphate (5’-P) terminus, olaparib consistently slowed PARP1 dissociation compared with the control without NAD+, whereas saruparib showed similar dissociation kinetics to the DMSO control (Fig. S5). The relatively faster PARP1 dissociation with saruparib compared to olaparib was not attributable to PAR-dependent PARP1 release, as detectable PARylation was absent under these conditions (Fig. 4F and Fig. S5C). Together, these findings demonstrate that conventional PARPis such as olaparib and talazoparib stabilize PARP1–DNA complexes beyond the effect of catalytic inhibition alone, whereas saruparib lacks this additional stabilization and forms more readily dissociable PARP1–DNA complexes.

### Low-nanomolar saruparib selectively kills HR-deficient cells

Having established that saruparib exhibits less stable PARP1 trapping than conventional PARPis, we examined its synthetic-lethal activity in HR-deficient cells. We employed TK6 cell lines in which BRCA1, BRCA2, or PALB2 was conditionally depleted using an auxin-inducible degron system (Fig. 5A). Depletion of BRCA1, BRCA2 or PALB2 markedly increased sensitivity to both olaparib and saruparib (Fig. 5B). Whereas olaparib-induced cytotoxicity progressively increased into the micromolar range, saruparib achieved near-maximal cytotoxicity at approximately 10 nM. Auxin treatment alone for 48 h induced detectable γH2AX, cleaved PARP1, and cleaved caspase-3 (0-h lanes). Concurrent treatment with auxin and either 10 μM olaparib or 10 nM saruparib for 24 or 48 h induced robust accumulation of these DNA damage and apoptotic markers (Fig. 5C). Thus, low-nanomolar saruparib is sufficient to induce robust DNA damage and apoptosis selectively in HR-deficient cells, with near-maximal cytotoxicity achieved at concentrations that fully inhibit PARP1 catalytic activity (∼10 nM, see Figure 1).

**Figure 5.**
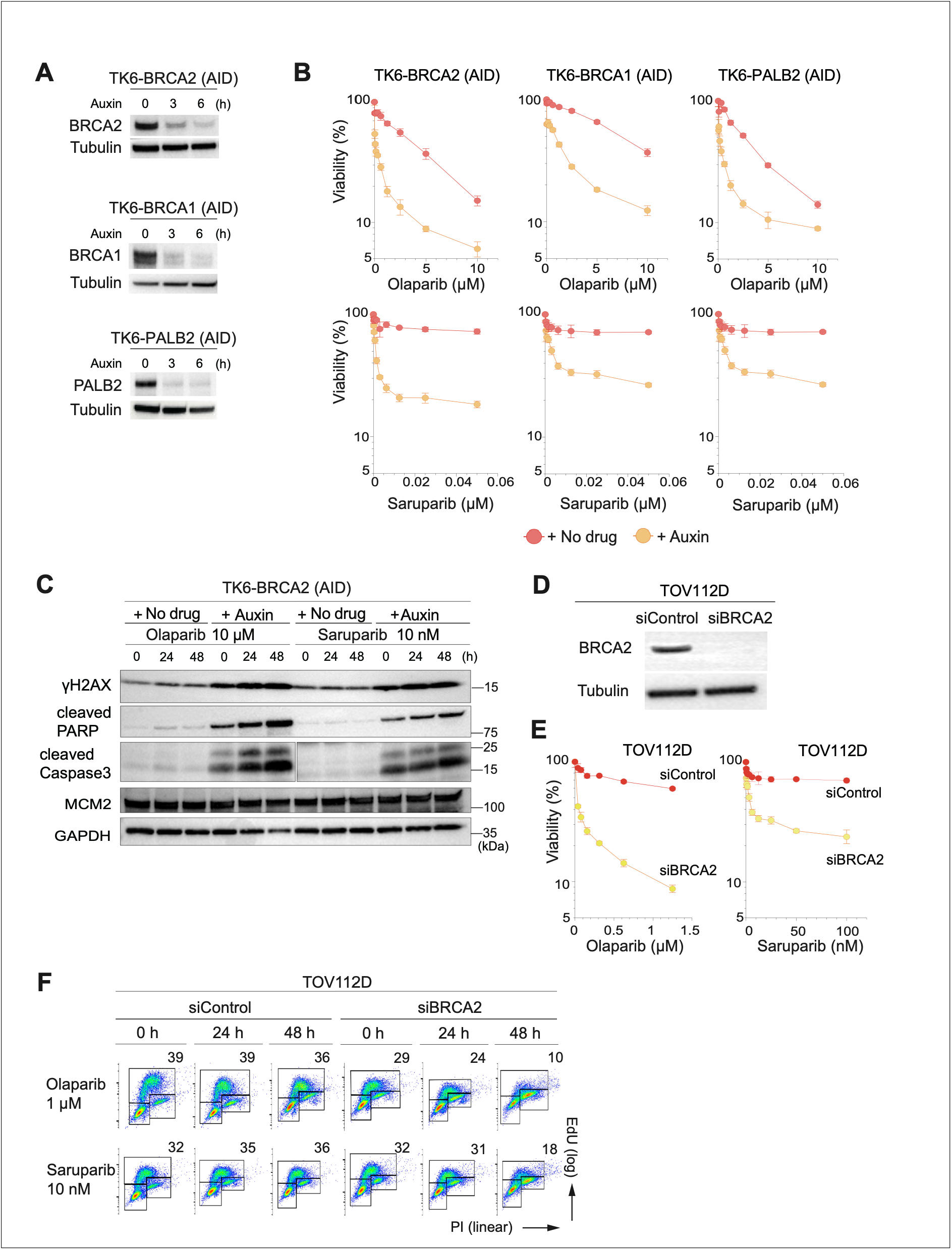
Low-nanomolar saruparib selectively kills HR-deficient cells. **A,** TK6 cells expressing auxin-inducible degron (AID)-tagged BRCA2, BRCA1, or PALB2 were treated with auxin (500 μM) for the indicated times. Protein depletion was confirmed by immunoblotting. **B,** Cell viability of TK6-BRCA2 (AID), TK6-BRCA1 (AID), and TK6-PALB2 (AID) cells treated with the indicated concentrations of olaparib or saruparib with or without auxin (500 μM) for 48 h. Viability was normalized to the corresponding untreated controls. Data are mean ± SD of triplicate wells and are representative of three independent experiments. **C** Time-course immunoblot analysis of γH2AX, cleaved PARP, and cleaved caspase-3 in TK6-BRCA2 (AID) cells treated with olaparib (10 μM) or saruparib (10 nM) with or without auxin (500 μM). For 0 h, cells were treated with auxin alone for 48 h; for 24 and 48 h, auxin and PARPi were co-administered for the indicated times. MCM2 and GAPDH were also analyzed. Data are representative of three independent experiments. **D,** Immunoblot analysis of BRCA2 in TOV112D cells transfected with control siRNA (siControl) or BRCA2-targeting siRNA (siBRCA2). **E,** Cell viability of TOV112D cells transfected with siControl or siBRCA2 and treated with the indicated concentrations of olaparib or saruparib. Data are presented as in **B**. **F,** Cell-cycle distribution of TOV112D cells transfected with siControl or siBRCA2 and treated with olaparib (1 μM) or saruparib (10 nM) for 24 or 48 h, analyzed as in Fig. 2E. Numbers indicate the percentage of cells in the replicating S-phase population. Data are representative of two independent experiments.

To validate these findings in an independent cancer cell model, we depleted BRCA2 using siRNA in TOV112D ovarian cancer cells (Fig. 5D). BRCA2 depletion increased sensitivity to both olaparib and saruparib, with saruparib again reaching near-maximal cytotoxicity at approximately 10 nM (Fig. 5E). Cell-cycle analysis showed that 1 μM olaparib markedly perturbed the S-phase population in BRCA2-depleted cells, while 10 nM saruparib produced a comparable perturbation selectively in BRCA2-depleted cells (Fig. 5F). Together, these results demonstrate potent synthetic-lethal activity of low-nanomolar saruparib in HR-deficient cells.

### PARPi sensitivity is determined by HR status and PARP-trapping properties rather than the extent of initial DNA damage

To further confirm the HR-dependent cytotoxicity of saruparib, we examined BRCA2-deficient PEO1 ovarian and Capan-1 pancreatic cancer cells and their PARPi-resistant derivatives carrying BRCA2 reversion mutations (26,27). BRCA2-revertant cells were resistant to talazoparib, olaparib, and saruparib (Fig. 6A, B), further confirming HR deficiency as a determinant of saruparib sensitivity.

**Figure 6.**
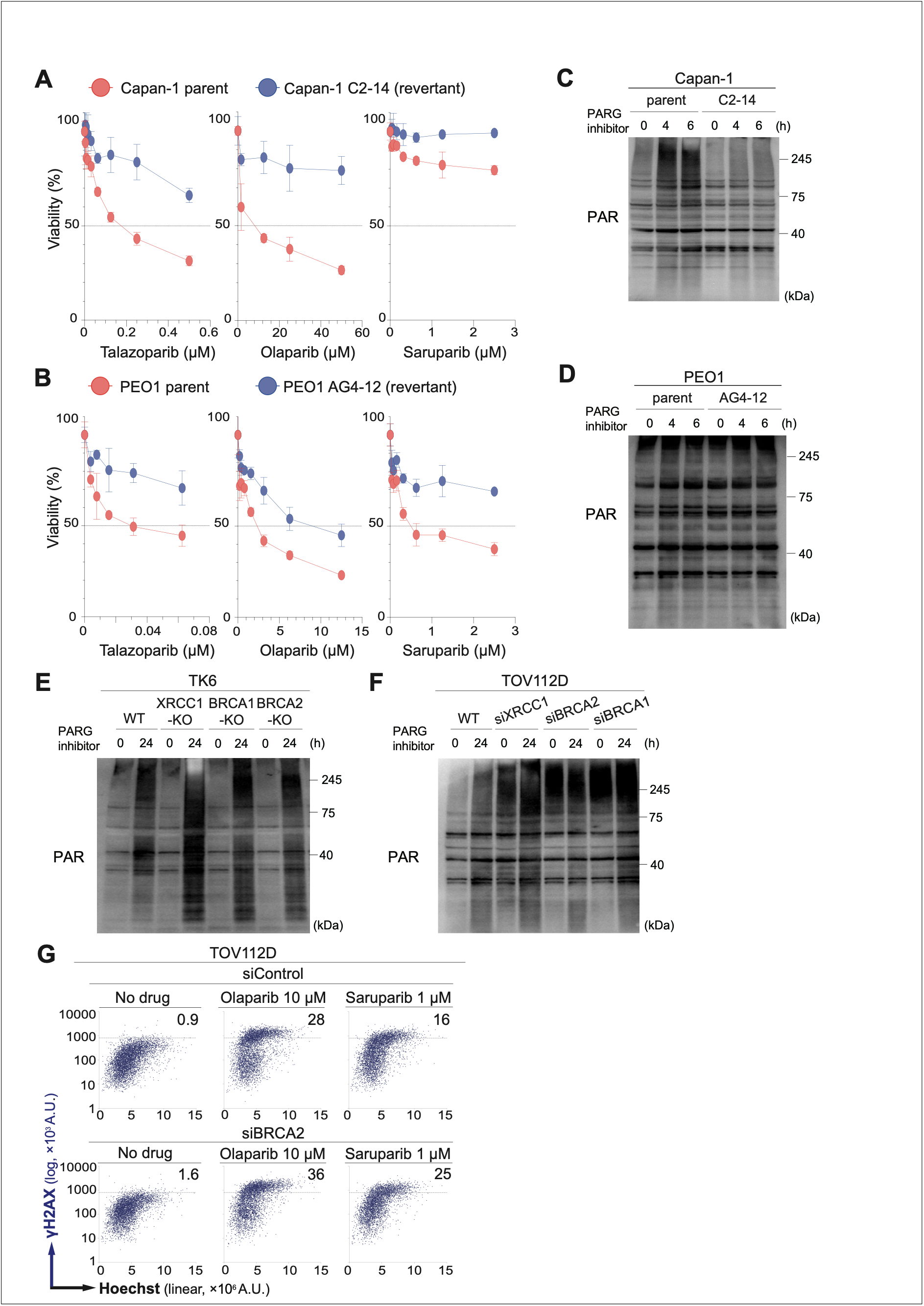
Saruparib induces replication-associated DNA damage throughout S phase. A, B, Cell viability of parental Capan-1 cells and the BRCA2-revertant C2-14 clone (A) and parental PEO1 cells and the BRCA2-revertant AG4-12 clone (B) treated with the indicated concentrations of PARP inhibitors for 48 h. Viability was normalized to the corresponding untreated controls.**C, D,** PAR accumulation in parental and BRCA2-revertant Capan-1 (C) and PEO1 (D) cells following treatment with PARG inhibitor (10 μM) for the indicated times, assessed by immunoblotting. **E, F,** PAR accumulation following PARG inhibition for 24 h in TK6 WT, XRCC1-KO, BRCA1-KD, and BRCA2-KD cells (E) and in TOV112D cells transfected with siControl, siXRCC1, siBRCA2, or siBRCA1 (). **G,** Single-cell analysis of γH2AX fluorescence intensity and DNA content in TOV112D cells transfected with siControl or siBRCA2 and treated with olaparib or saruparib for 4 h. γH2AX fluorescence intensity was measured in the Alexa Fluor 488 channel, and DNA content was assessed by Hoechst fluorescence intensity. Numbers in each panel indicate the percentage of cells with a γH2AX fluorescence intensity ≥ 800,000 A.U.

Given the relatively weak and unstable PARP1 trapping induced by saruparib, we next sought to understand how saruparib selectively kills HR-deficient cells. Previous studies have shown that endogenous PARP activity is elevated in BRCA2-deficient and other HR-deficient cells (43). Consistent with these findings, PARG inhibition resulted in greater PAR accumulation in BRCA2-deficient PEO1 and Capan-1 cells than in their BRCA2-revertant derivatives (Fig. 6C, D). Increased PAR accumulation was also observed following BRCA1 or BRCA2 depletion in TK6 and TOV112D cells (Fig. 6E, F). These findings suggest that HR-deficient cells augment PARP1 engagement at endogenous DNA lesions or replication-associated DNA intermediates.

To determine whether the extent and cell-cycle distribution of the early DNA damage response induced by PARPis differ according to HR status or the trapping properties of individual inhibitors, we performed high-content imaging of γH2AX in control and BRCA2-depleted cells treated with olaparib or saruparib. Cells with a γH2AX intensity of zero, which were predominantly distributed in the G1/sub-G1 population (Fig. S6), were excluded from the plots in Fig. 6G because zero values cannot be displayed on the logarithmic scale. A 4-h treatment with either olaparib or saruparib induced γH2AX throughout S phase in both control and BRCA2-depleted cells (Fig. 6G). Notably, despite the increased PARP activity observed in BRCA2-deficient cells (Fig. 6C–F), BRCA2 depletion did not substantially increase γH2AX levels following PARPi treatment. Thus, modest differences in endogenous PARP activity do not appear to be the primary determinant of PARPi-induced cytotoxicity. Rather, together with the marked differences in cell viability, these findings indicate that cellular sensitivity is governed predominantly by HR status and by the trapping properties of individual PARPis, which together determine the fate of replication-associated DNA lesions.

## Discussion

In this study, we investigated the pharmacological properties of the PARP1-selective inhibitor saruparib across multiple genetic and DNA damage settings. A consistent feature of saruparib is the limited dose dependency of its cytotoxicity above approximately 10 nM (Figs. 2, 3, 5, and 6), a concentration range that also corresponded to near-maximal inhibition of cellular PARylation (Fig. 1). Although saruparib induces detectable PARP1 trapping, its trapping properties appear distinct from those of conventional PARPis such as olaparib and talazoparib in terms of both complex stability and trapping-associated cytotoxicity (Figs. 3 and 4). Overall, our findings suggest that saruparib closely approximates the original concept of PARP–BRCA synthetic lethality proposed in 2005, achieving potent cytotoxicity in HR-deficient cells while largely sparing HR-proficient cells (1,2).

Whether these distinctive properties of saruparib reflects its PARP1 selectivity or its specific interaction with PARP1 remains unresolved. However, the lack of PARP2 inhibition alone is unlikely to explain its limited trapping-associated cytotoxicity. Niraparib has relatively weak activity against PARP2 but exhibits potent PARP1 trapping (Fig. 1B) (5,14). Conventional PARPis retain potent trapping and trapping-associated cytotoxicity in chicken DT40 cells, which naturally lack PARP2 (5,6,44). These observations suggest that the distinctive behavior of saruparib arises from its specific interaction with PARP1 rather than simply from PARP1 selectivity. Direct comparison with other PARP1-selective inhibitors should clarify whether readily reversible PARP1–DNA complexes are a general property of PARP1-selective inhibition or a feature specific to saruparib.

Our biochemical measurements further support the notion that saruparib modulates PARP1– DNA interactions differently from conventional PARPis. Whereas olaparib and talazoparib stabilize PARP1–DNA complexes beyond the stability observed in the absence of NAD⁺, saruparib does not confer additional stabilization and shows features consistent with more readily reversible PARP1–DNA binding. These findings are consistent with an allosteric model in which inhibitor binding at the catalytic domain modulates PARP1–DNA interactions through long-range conformational changes (5,45). Conventional cellular trapping assays based on immunofluorescence or immunoblotting measure the amount of chromatin-associated PARP1 but do not readily distinguish stable trapping from transient or less stable PARP1–DNA binding. Our cell-free assay may therefore provide a complementary approach for evaluating inhibitor-induced stabilization or destabilization of PARP1–DNA complexes, including potential allosteric trapping and detrapping effects.

The limited stability of saruparib-induced PARP1–DNA complexes may explain its distinct response to MMS (Fig. 3). If saruparib-bound PARP1 dissociates relatively readily upon collision with an advancing replication fork, most encounters may be resolved without severe DNA damage. However, the large increase in PARP1-binding lesions induced by MMS would increase such encounters and the probability of generating severe replication-associated lesions and trapping-dependent cytotoxicity. This model provides a potential explanation for why saruparib cytotoxicity plateaus once PARP1 is fully inhibited, rather than progressively increasing with drug concentration as observed with conventional PARPis.

An important question raised by our findings is how saruparib achieves potent synthetic lethality in HR-deficient cells despite forming relatively unstable PARP1–DNA complexes. A recent genetic study identified the PARP1 L777P mutation in cells that had acquired resistance to saruparib and used this mutant to examine the relationship between catalytic inhibition, PARP1 trapping, and cytotoxicity (20). For conventional PARPis such as olaparib and veliparib, L777P reduced catalytic inhibition but paradoxically increased PARP1 trapping and further sensitized BRCA-deficient cells, providing strong genetic evidence that enhanced trapping can directly increase cytotoxicity in the HR-deficient setting. In contrast, saruparib behaved differently: the same L777P mutation impaired both its catalytic inhibition and its ability to trap PARP1. Thus, in the case of saruparib, this mutant does not separate catalytic inhibition from trapping and therefore cannot establish which activity is responsible for its HR-selective cytotoxicity. Importantly, our present data demonstrate that saruparib forms PARP1–DNA complexes with properties distinct from those induced by conventional PARPis, including weaker and less stable trapping. Accordingly, the relationship between trapping and cytotoxicity defined for conventional PARPis should not be assumed to apply directly to saruparib.

HR-deficient cells exhibit elevated endogenous PARP1 activity (Fig 6), potentially reflecting an increased burden of endogenous PARP1-engaging lesions. However, PARP1 activation in HR-deficient cells was substantially lower than that in XRCC1-deficient cells (Fig. 6E, F), which remained relatively insensitive to saruparib, indicating that increased PARP1 activation alone cannot account for saruparib cytotoxicity.

Recent studies have proposed a trapping-independent mechanism of PARPis synthetic lethality involving transcription–replication conflicts that preferentially arise during early S phase (19). Our finding that saruparib-induced γH2AX in BRCA2-deficient cells was observed throughout S phase rather than being restricted to early S phase (Fig. 6G) suggests that its synthetic-lethal activity is not limited to this mechanism. Instead, our results support a broader model in which endogenous PARP1-engaging lesions persist following catalytic PARP1 inhibition and are converted into replication-associated DNA damage throughout S phase. The inability of HR-deficient cells to repair these lesions may therefore explain why saruparib retains potent synthetic-lethal activity despite its relatively unstable PARP1 trapping.

### Clinical implications

Dosing of conventional PARPis has been established at exposures substantially exceeding those required for catalytic PARP inhibition, consistent with their concentration-dependent trapping-associated cytotoxicity (4). In contrast, saruparib showed little additional cytotoxicity above approximately 10 nM across multiple experimental settings once near-maximal catalytic inhibition was achieved. These findings raise the possibility that optimal saruparib exposure could be defined primarily by sustained catalytic PARP1 inhibition. Integration of these mechanistic findings with clinical pharmacokinetic and pharmacodynamic data will be important to determine how currently used doses, including 60 mg/day, relate to the exposure required for sustained PARP1 inhibition.

An important clinical question is how broadly the synthetic-lethal activity of saruparib extends beyond BRCA1/2 deficiency. The concept of “BRCAness” encompasses defects in multiple HR and DNA damage response genes that can confer sensitivity to conventional PARPis, including RAD51, FANCA, CDK12, ATR, and ATM (34,35,46,47). However, sensitivity to conventional PARPis may reflect varying contributions of catalytic inhibition and PARP trapping. Given the relatively limited trapping-associated cytotoxicity of saruparib, it remains to be determined whether these broader forms of BRCAness cause sufficient impairment of HR-mediated repair to confer sensitivity to saruparib. Thus, rather than the presence of individual DNA repair gene alterations per se, the functional capacity for HR-mediated repair may be a critical determinant of saruparib sensitivity.

The emergence of saruparib as a next-generation PARP inhibitor may broaden the potential applications of PARP inhibition. Saruparib could, in principle, eliminate HR-deficient cancer cells while imposing substantially less genotoxic stress on surrounding HR-proficient cells, raising the possibility of cancer treatment or prevention in genetically defined high-risk populations, such as individuals with hereditary breast and ovarian cancer (HBOC). However, genetic studies have shown that PARP1 loss can critically influence cellular outcome, with partial PARP1 reduction or inhibition preceding BRCA2 loss permitting synthetic viability under specific experimental conditions (48). Thus, the long-term consequences of sustained PARP1 inhibition on the emergence and fate of HR-deficient cells require careful investigations. Nevertheless, two decades after the discovery of PARP–BRCA synthetic lethality, saruparib may represent a pharmacological realization of its original principle: potent and selective elimination of HR-deficient cells with minimal cytotoxicity in HR-proficient cells. This property may provide a foundation for a more precise mechanism-guided use of PARP inhibition in cancer therapy.

## Author contributions

Conceptualization: S.N., Y.P., J.M.

Resources: T.I., S.I., S.D., K.H., H.S., T.S., K.S., J.M.

Data curation: S.N., J.M.

Formal analysis: S.N., J.M.

Supervision: T.S., K.S., J.M.

Funding acquisition: T.S., K.S., J.M.

Investigation: S.N., H.O., R.K., E.K., K.U., T.I., S.I., J.M.

Visualization: S.N., J.M.

Methodology: S.N., S.D., H.S., J.M.

Writing – original draft: S.N., Y.P., J.M.

Project administration : J.M.

Writing – review & editing: All authors

## Acknowledgement

We thank Mr. Mikio Nakane for technical assistance and Prof. Toshiyasu Taniguchi (Tokai University School of Medicine) for providing the Capan-1 and PEO1 cell lines and their BRCA2-revertant derivatives. We also thank the TK6 Mutants Consortium (http://www.nihs.go.jp/dgm/tk6.html) for their critical help in establishing methods for gene-disruption using the TK6 cell line. We acknowledge the support of ADRES (Advanced Research Support Center, Ehime University).

## Financial support

This work was supported in part by the Japan Society for the Promotion of Science, Grants-in-Aid for Scientific Research (23K27459 and 26K02006 to J.M.); the Japan Science and Technology Agency (JST), grant number JPMJFR2056 (JST FOREST Program to J.M.); the Naito Foundation (to J.M.); the Uehara Memorial Foundation (to J.M.); the Takeda Science Foundation (to J.M.); and research funds from donations to the Department of Obstetrics and Gynecology, Ehime University (to T.S. and K.S.).

## Conflict of interest disclosure statement

J. Murai has received lecture fees from Takeda Pharmaceutical Company, Pfizer, and AstraZeneca. All other authors declare no conflicts of interest.

