## Supplementary material for "Saruparib recapitulates PARP–BRCA synthetic lethality with PARP1-trapping properties limiting its cytotoxicity": GA and Supplementary figures

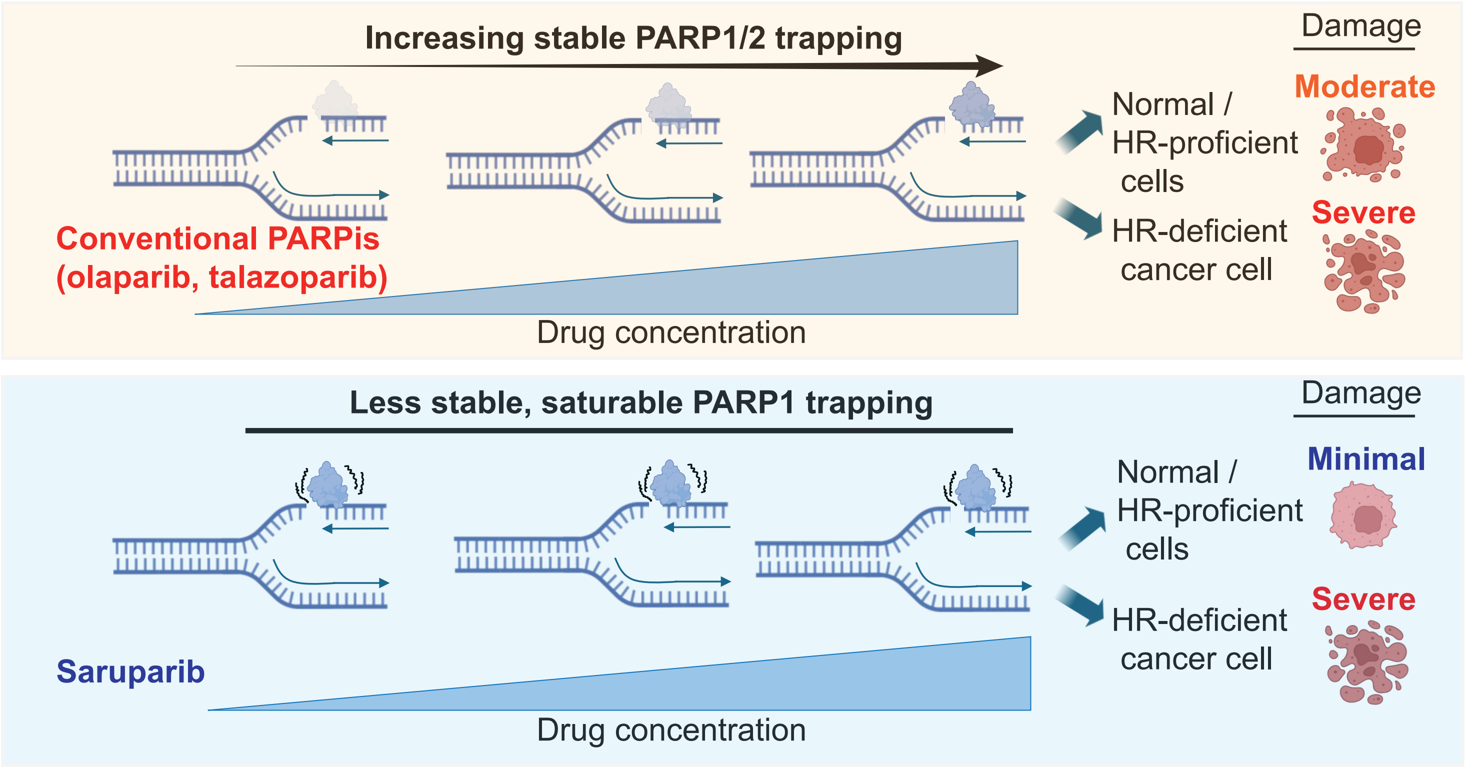
Increasing stable PARP1/2 trapping
Damage
Normal / HR-proficient cells
Moderate
HR-deficient cancer cell
Severe
Conventional PARPis (olaparib, talazoparib)
Drug concentration
Less stable, saturable PARP1 trapping
Damage
Normal / HR-proficient cells
Minimal
HR-deficient cancer cell
Severe
Saruparib
Drug concentration

**Supplementary Figures**

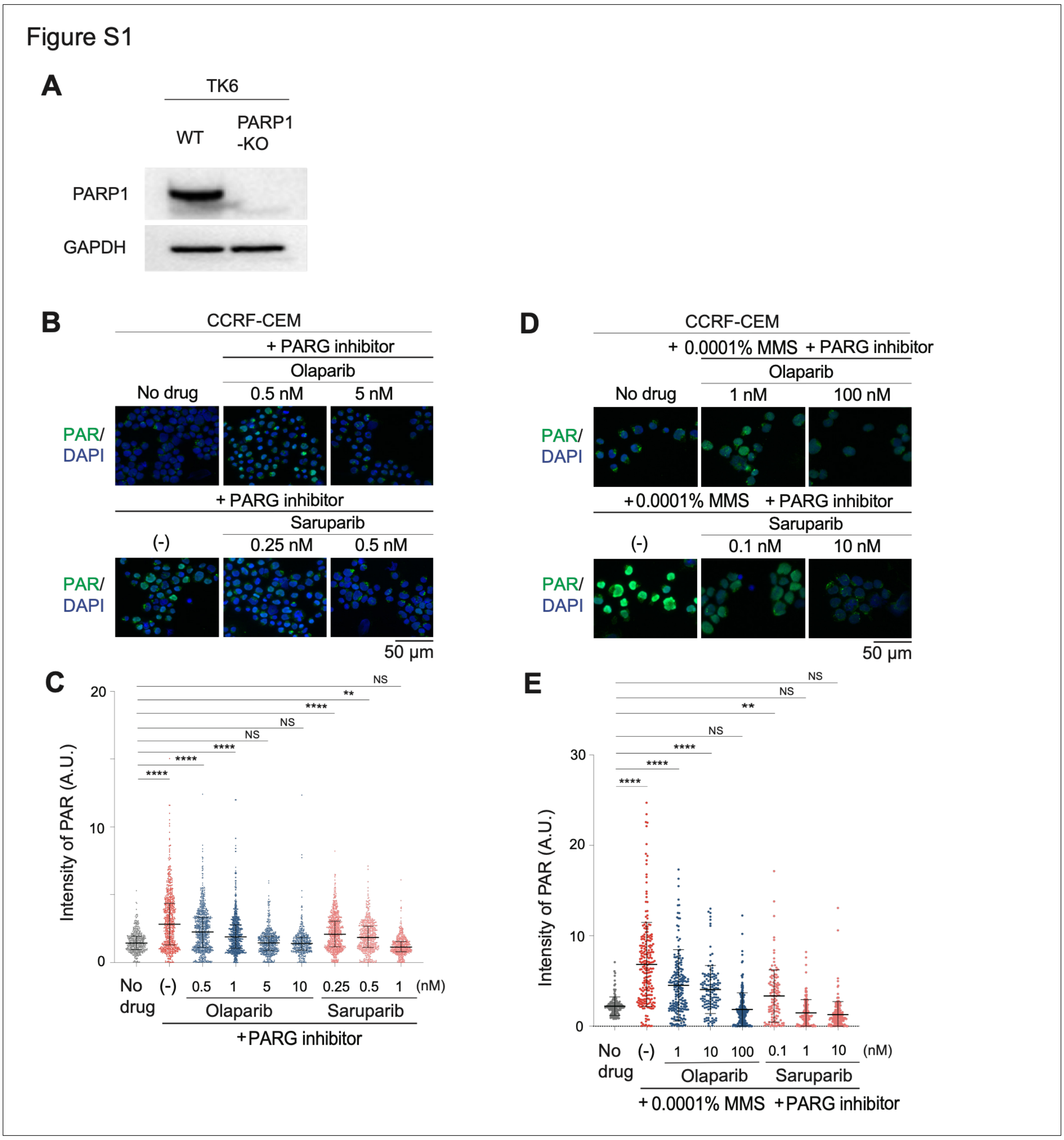


**Figure S1. Saruparib potently inhibits PARP1 catalytic activity in CCRF-CEM cells.**

**A,** Immunoblot analysis of PARP1 expression in TK6 WT and PARP1-KO cells. GAPDH served as a loading control. **B, C,** Representative images (**B**) and quantification (**C**) of PAR immunofluorescence in CCRF-CEM cells treated with the indicated concentrations of olaparib or saruparib in the presence of PARG inhibitor (10 µM) for 4 h, analyzed as in Fig. 1D and 1E. **D, E,** Representative images (**D**) and quantification (**E**) of PAR immunofluorescence in CCRF-CEM cells treated with the indicated concentrations of olaparib or saruparib in combination with MMS (0.0001%) and PARG inhibitor (10 µM) for 4 h, analyzed as in Fig. 1D and 1F. For **C** and **E**, each dot represents an individual cell; bars indicate mean ± SD (C; n = 411–963,

E; n = 122–230 cells per condition). NS, not significant; **P < 0.01; ****P < 0.0001 by one-way ANOVA followed by Tukey’s multiple-comparisons test.

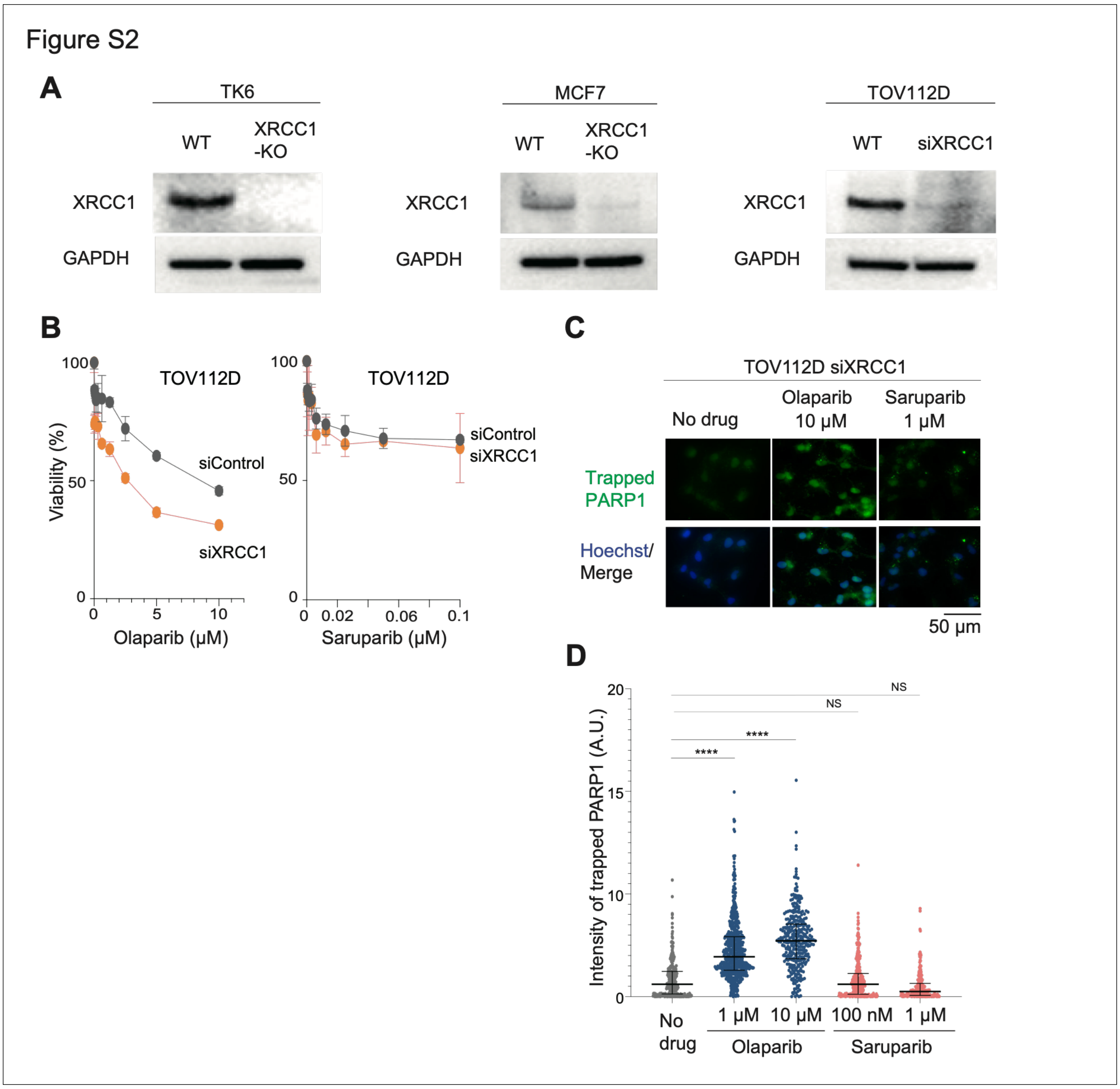


**Figure S2. Saruparib exhibits minimal PARP1 trapping even in XRCC1-deficient cells.**

**A,** Immunoblot analysis of XRCC1 expression in TK6 WT and XRCC1-KO cells, MCF7 WT and XRCC1-KO cells, and TOV112D cells transfected with siXRCC1. GAPDH served as a loading control. **B,** Cell viability of TOV112D cells transfected with siControl or siXRCC1 and treated with the indicated concentrations of olaparib or saruparib for 48 h. Viability was normalized to the corresponding untreated controls. Data are mean ± SD of technical replicates and are representative of independent experiments. **C,** Representative images of chromatin-bound PARP1 in TOV112D cells depleted of XRCC1 and treated with olaparib or saruparib, assessed as in Fig. 2C. Scale bar, 50 μm. **D,** Quantification of chromatin-bound PARP1 fluorescence intensity in TOV112D cells depleted of XRCC1 and treated with the indicated concentrations of olaparib or saruparib. Each dot represents an individual cell; bars indicate mean ± SD (n = 232–618 cells per condition). NS, not significant; ****P < 0.0001 by one-way ANOVA followed by Tukey's multiple-comparisons test.

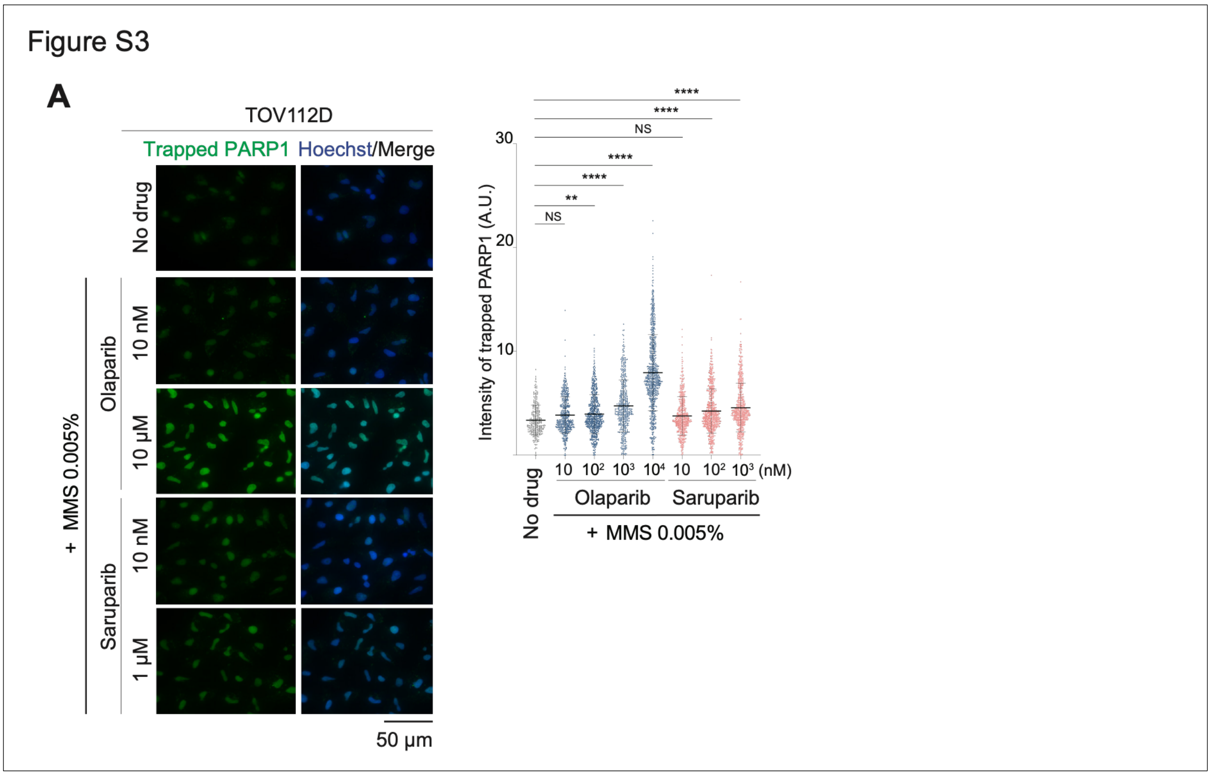


**Figure S3. Saruparib induces saturable PARP1 trapping in the presence of MMS.**

**A,** Representative images and quantification of chromatin-bound PARP1 in TOV112D cells treated with the indicated concentrations of olaparib or saruparib in combination with MMS (0.005%), assessed as in Fig. 3C and 3D. Each dot represents an individual cell; bars indicate mean ± SD (n = 320–832 cells per condition). NS, not significant; $**P < 0.01$; $****P < 0.0001$ by one-way ANOVA followed by Tukey's multiple-comparisons test. Scale bar, 50 µm.

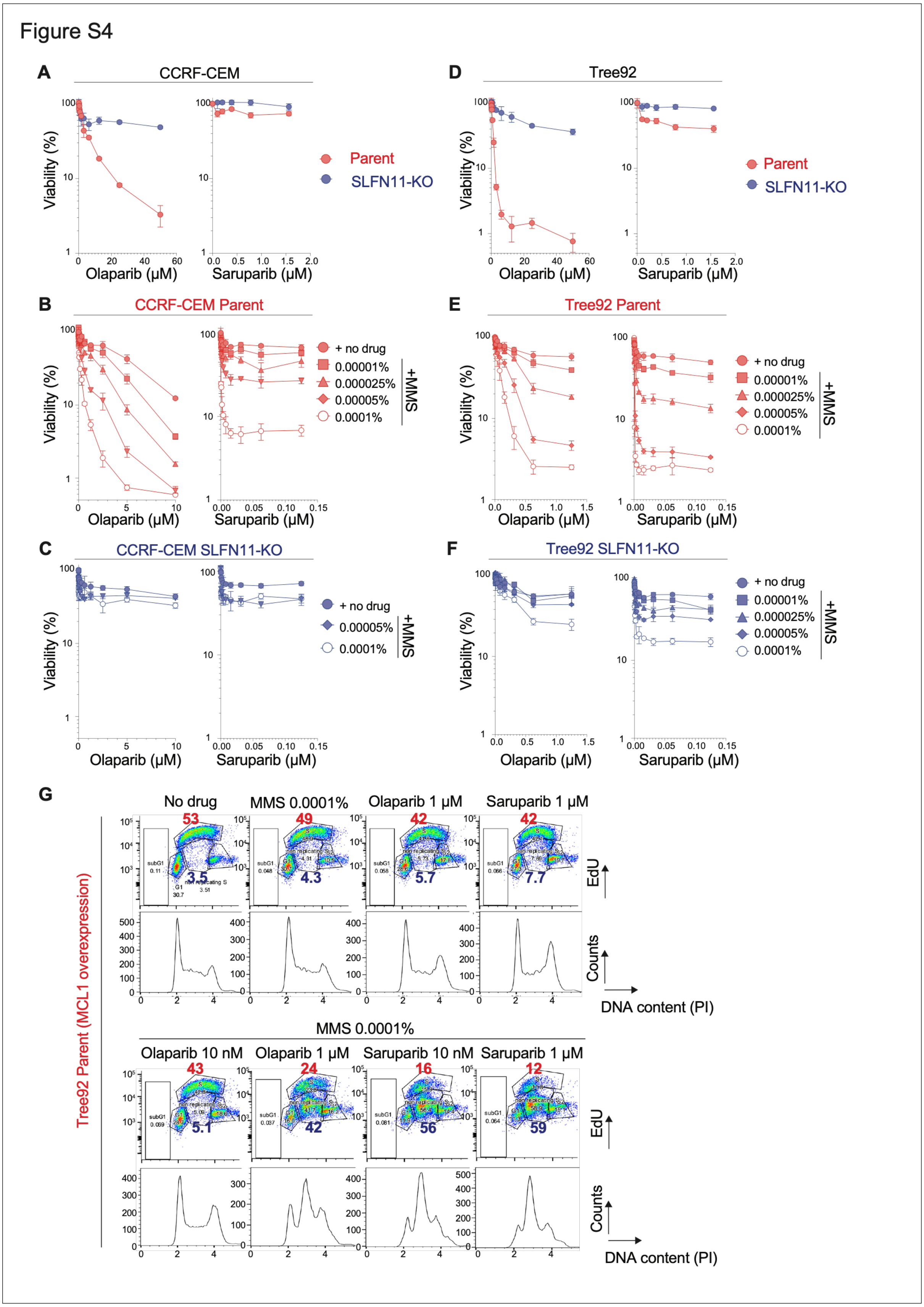


**Figure S4. Saruparib and olaparib induce replication stress and SLFN11-dependent cytotoxicity in the presence of MMS.**

**A, D,** Cell viability of parental and SLFN11-KO CCRF-CEM (A) and Tree92 (D) cells treated with the indicated concentrations of olaparib or saruparib for 96 h. **B, C, E, F,** Cell viability of parental (B, E) and SLFN11-KO (C, F) CCRF-CEM (B, C) and Tree92 (E, F) cells treated with the indicated concentrations of olaparib or saruparib alone or in combination with MMS for 96 h. Viability was normalized to the corresponding untreated controls. Data are mean ± SD of technical replicates and are representative of two independent experiments. **G,** Cell-cycle distribution of MCL1-overexpressing parental Tree92 cells treated with MMS, olaparib, or saruparib alone or with the indicated concentrations of olaparib or saruparib in combination with MMS for 24 h. Cells were analyzed by EdU incorporation and propidium iodide staining as in Fig. 2E. Red numbers indicate the percentages of replicating S-phase cells, while blue numbers indicate the percentages of unreplicating S-phase cells.

Figure S5

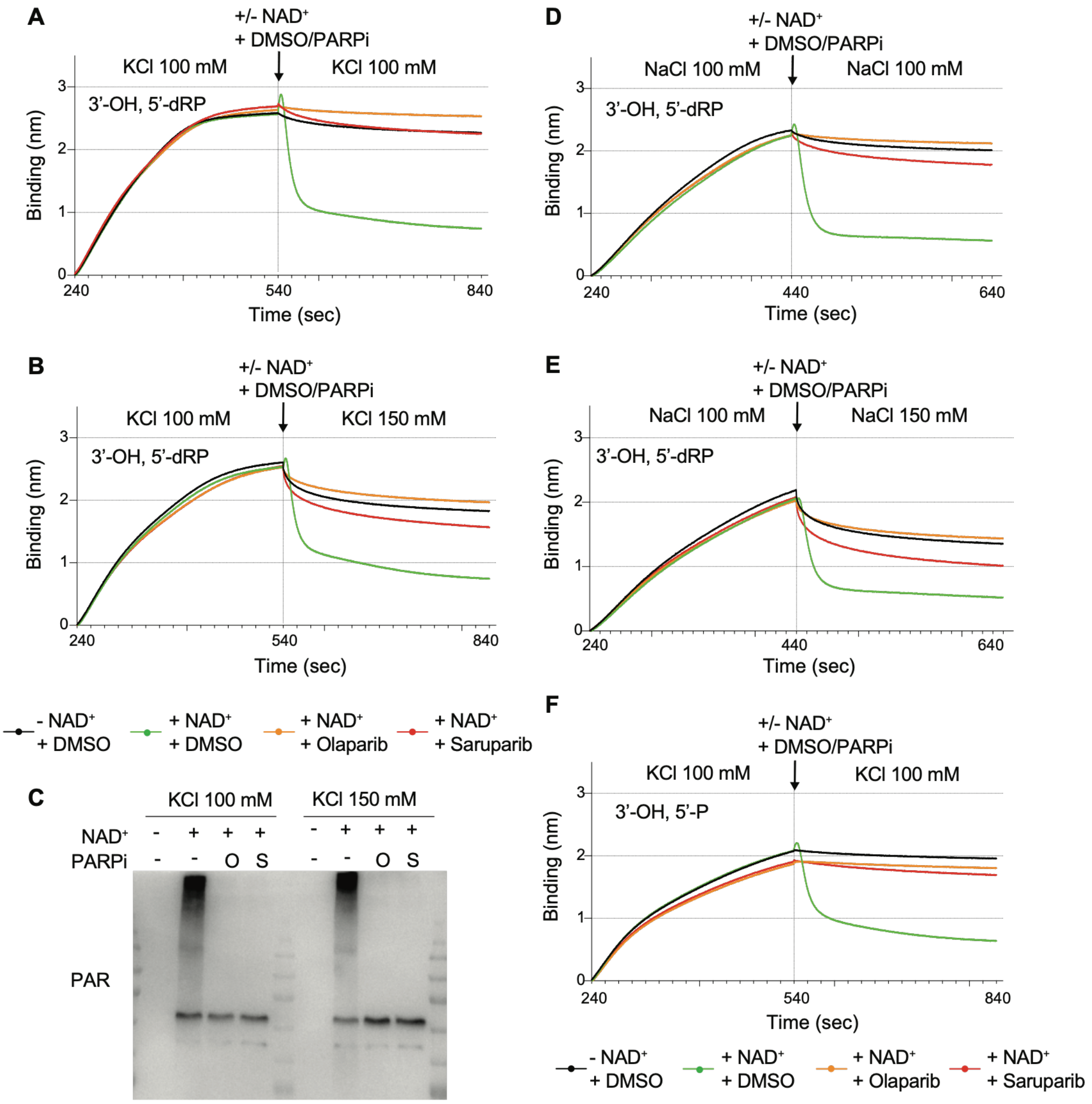


**G**

| Substrate | Buffer | - NAD+ + DMSO kdis (1/s) | + NAD+ + DMSO kdis (1/s) | + NAD+ + Olaparib kdis (1/s) | + NAD+ + Saruparib kdis (1/s) |
|---|---|---|---|---|---|
| 3'-OH, 5'-dRP | KCl 100 mM | 1.097E-02 | 4.680E-02 | 7.530E-03 | 1.055E-02 |
| 3'-OH, 5'-dRP | KCl 150 mM | 1.434E-02 | 3.674E-02 | 1.107E-02 | 1.245E-02 |
| 3'-OH, 5'-dRP | NaCl 100 mM | 1.726E-02 | 7.712E-02 | 1.254E-02 | 1.629E-02 |
| 3'-OH, 5'-dRP | NaCl 150 mM | 2.100E-02 | 7.575E-02 | 1.815E-02 | 2.021E-02 |
| 3'-OH, 5'-P | KCl 100 mM | 6.038E-03 | 3.212E-02 | 2.528E-03 | 5.033E-03 |

**Figure S5. Saruparib forms less stable PARP1–DNA complexes than olaparib.**

**A, B,** Representative BLI sensorgrams showing PARP1 dissociation from the 3′-OH/5′-dRP DNA substrate in 100 mM (**A**) or 150 mM (**B**) KCl, analyzed as in Fig. 4D and 4E. 10 μM olaparib or 10 μM saruparib was used. **C,** Dissociation solutions from **A** and **B** were analyzed by immunoblotting for PAR to confirm PARP1 catalytic activity and its inhibition by olaparib (O) or saruparib (S). **D, E,** BLI analysis performed as in **A** and **B**, except that KCl was replaced with

100 mM (**D**) or 150 mM (**E**) NaCl. **F,** BLI analysis of PARP1 dissociation from a 3′-OH/5′-phosphate DNA substrate in 100 mM KCl, performed as in **A**. **G,** Dissociation rate constants (k_dis) calculated from the BLI curves under the indicated DNA substrate and ionic conditions.

Figure S6

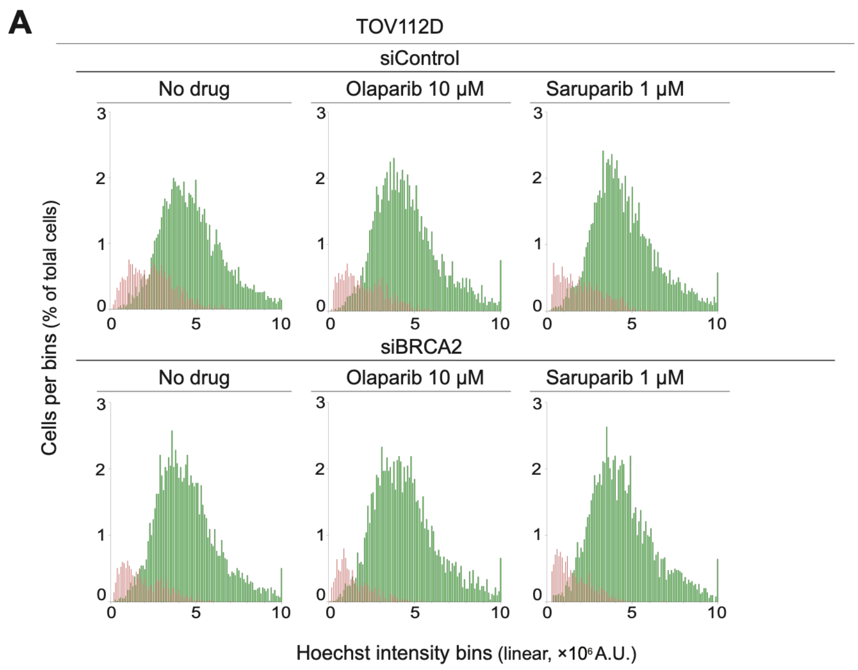


**Figure S6. Distribution of γH2AX-positive and -negative cells according to DNA content in TOV112D cells.**

**A,** Distribution of the TOV112D cell populations used for the analysis in Fig. 6G. Cells were transfected with siControl or siBRCA2 and treated with olaparib or saruparib for 4 h. Hoechst fluorescence intensity was binned at intervals of 100,000 A.U. The y-axis represents the percentage of cells in each bin relative to the total number of analyzed cells. Cells with no detectable γH2AX signal are shown in orange, whereas cells with detectable γH2AX signal are shown in green. The γH2AX-positive population shown in green was used for the analysis in Fig. 6G.
